# Gas6 drives Zika virus-induced neurological complications in humans and congenital syndrome in immunocompetent mice

**DOI:** 10.1101/2021.01.27.428402

**Authors:** Joao Luiz Silva-Filho, Lilian Gomes de Oliveira, Leticia Monteiro, Pierina L. Parise, Nagela G. Zanluqui, Carolina M. Polonio, Carla Longo de Freitas, Daniel A. Toledo-Teixeira, William M. Souza, Najara Bittencourt, Mariene R. Amorim, Julia Forato, Stéfanie Primon Muraro, Gabriela Fabiano de Souza, Matheus Cavalheiro Martini, Karina Bispos-dos-Santos, Carla C. Judice, Maria Laura Costa, Rodrigo N. Angerami, André R. R. Freitas, Mariangela R. Resende, Márcia T. Garcia, Maria Luiza Moretti, The Zika-Unicamp Network, Laurent Renia, Lisa F. P. Ng, Carla V. Rothlin, Fabio TM Costa, Jean Pierre Schatzmann Peron, José Luiz Proença-Modena

## Abstract

Zika virus (ZIKV) has the ability to cross placental and brain barriers, causing congenital malformations in neonates and neurological disorders in adults. However, the pathogenic mechanisms of ZIKV-induced neurological complications in adults and congenital malformations remain unknown. Gas6 is a soluble TAM receptor ligand able to promote flavivirus internalization and downregulation of immune responses. Here we demonstrate high Gas6 levels in the serum of patients with neurological complications which correlated with downregulation of genes associated with the type I IFN responses as consequence of *Socs1* upregulation. Gas6 gamma-carboxylation is essential for ZIKV replication in monocytes, the main source of this protein. Gas6 also facilitates ZIKV replication in adult immunocompetent mice enabled susceptibility to transplacental infection and congenital malformations. Our data thus indicate that ZIKV promotes the upregulation of its ligand Gas6, which contributes to viral infectivity and drives the development of severe adverse outcomes during ZIKV infection.

## INTRODUCTION

In 2016, Zika virus (ZIKV) emerged as an important global health problem, starting in South America, and then spreading to more than 94 countries worldwide. First discovered in 1947 in Uganda, Africa, it has not been considered a threat to human health, until the outbreaks in Yap, Micronesia (2007) and French Polynesia (2013) [1]. Most individuals are asymptomatic or develop a benign febrile disease characterized by cutaneous rash and conjunctivitis. However, it was further shown that ZIKV can cross the placental barrier and reach foetal tissues causing the congenital ZIKV syndrome (CZS), that may range from foetal growth restriction and microcephaly to severe retinal damage and arthrogryposis [1–3]. This was unprecedented and demanded great efforts of the scientific community to understand the underlying mechanisms involved in host-virus interaction, mainly those related to susceptibility and pathogenicity.

Central nervous system (CNS) manifestations after congenital infection, such as brain calcifications, lissencephaly, ventricular hypertrophy and microcephaly, are among the most notable and concerning outcomes of ZIKV infection in newborns [4–6]. However, severe ZIKV infection is not limited to newborns: neurological manifestations as acute myelitis, encephalitis, meningoencephalitis and Guillain-Barré syndrome can also occur in adults [7–9]. This raises not only the question on what are the genes that confer susceptibility to ZIKV neuropathology, but also, what are the cellular and molecular mechanisms orchestrating such phenomenon. In this context, we believe the interaction between viral particles with host cells, the first step of infection, may be of pivotal relevance. This interaction dictates more than the viral tissue tropism, but also triggers a diversity of intracellular pathways that may greatly account for either failure or success of infection [10, 11].

Growth arrest-specific 6 (Gas6) is a 75 KDa secreted protein composed of an N-terminal Gla domain, followed by four epidermal growth factor (EGF)-like domains and a C-terminal SHBG domain [12]. Upon γ-carboxylation of the Gla domain, Gas6 is able to interact with TAM (Tyro3, Axl and Mer) receptors and phosphatidylserine (PtdSer) promoting phagocytic internalization of the apoptotic bodies [12]. TAM activation triggers different signalling pathways involved in cell survival, mainly orchestrated by PI3K and Akt [12, 13]. Interestingly, PtdSer containing viruses, such as flaviruses and filoviruses, may also bind to Gas6, activating clathrin-mediated phagocytic internalization and subverting cellular immune response by activating negative regulators of anti-viral cytokines, as SOCS-1 and SOCS-3 [10, 11, 14], negatively regulating type I interferon receptor (IFNAR) signalling pathway [12, 13–18]. However, the overall interplay of viral and host factors, such as GAS6, to orchestrate the clinical course of ZIKV infection is not fully understood.

Here we investigated the role of the PtdSer ligand Gas6 in the pathogenesis of ZIKV infection. We observed that circulating levels of Gas6 are upregulated in the serum of ZIKV- infected patients, including pregnant women, and a further increase occurs in patients with neurological complications. Concurrently, there is a reduced transcriptional expression of genes associated with type I IFN responses and other immune signatures, probably as consequence of *Socs1* upregulation in peripheral blood cells. ZIKV infected monocytes were an important source of Gas6 production and Gas6 gamma-carboxylation was essential for ZIKV replication. Conversely, we also demonstrate that Gas6 is upregulated in ZIKV-infected pregnant mice and that pre-incubation of ZIKV with recombinant Gas6 facilitates ZIKV replication and rendered them susceptible to transplacental infection and congenital malformations. We thus propose that Gas6 dampens antiviral immune response in the periphery, promoting viral replication and facilitating severe clinical outcome. Collectively, our data brings novelty to the role of Gas6 on the understanding of ZIKV pathogenesis, as a relevant host factor driving severe outcomes, such as neurological complications. Addressing this issue could help develop predictive approaches for early diagnostics and open new possibilities to develop effective treatment against severe complications.

## RESULTS

### Patients

In this cross-sectional study, peripheral blood, serum, and urine samples were collected from 90 patients and 13 healthy donors during ZIKV epidemics in Brazil. These samples were originally collected between February 2016 and June 2017 in different hospitals in the city of Campinas. Included participants were recruited based on their clinical symptoms during hospital admission and according to the results of ZIKV laboratorial tests [9]. In brief, we included all patients with neurological complications suspected of arbovirus infection that we had access to during the period (inclusion and exclusion criteria are described in star methods section). In addition, we included samples from 57 patients with a mild disease, all positive for ZIKV by RT-qPCR (Non-Neuro^ZIKV^). Of the 33 patients with neurological complications, 19 (60%) were positive for ZIKV in previous tests or during hospitalization (Neuro^ZIKV^) and 14 were negative for ZIKV (Neuro^NON-ZIKV^). Neurological complications started between 2 and 15 days after onset of acute symptoms (mean of 4 days) in ZIKV patients. At the day of sample collection, there was no difference in ZIKV RNA load in the peripheral blood samples from the Non-Neuro^ZIKV^ and Neuro^ZIKV^ patients (median near of 2000 copies/mL of the RNA viral in both groups of patients) (Table 1 summarizes the demographic data and clinical manifestations).

**Table 1.** Sociodemographic, clinical and laboratorial findings in the considered groups included in this study

| Characteristic | Patients No (%) |  |  |  |
| --- | --- | --- | --- | --- |
|  | Non-Neuro<sup>ZIKV</sup><br>(n=57) | Neuro<sup>ZIKV</sup><br>(n=19) | Neuro<sup>NON-ZIKV</sup><br>(n=14) | HD<br>(n=13) |
| **Demographics** |  |  |  |  |
| Age, median (range) in years | 31 (1 – 69) | 32 (1 – 82) | 26 (1 – 74) | 29 (18 – 45) |
| Female sex | 42 (73.7) | 11 (57.8) | 6 (42.8) | 8 (61.5) |
| **Preceding symptoms** |  |  |  |  |
| Fever | 35 (61.5) | 10 (52.6) | 7 (50.0) | NA |
| Rash | 37 (65) | 8 (42.1) | 4 (28.5) | NA |
| Conjunctivitis | 24 (42.1) | 7 (36.8) | 2 (14.3) | NA |
| Headache | 22 (38.6%) | 5 (26.3) | 3 (21.4) | NA |
| Arthralgias | 12 (21%) | 3 (15.7) | 3 (21.4) | NA |
| Myalgias | 19 (33.3%) | 2 (10.5) | 1 (7.2) | NA |
| Time from onset of viral symptoms to ZIKV diagnosis, median (range) in days | 3 (0 – 10) | 4 (1 – 10) | NA | NA |
| **Laboratory tests** |  |  |  |  |
| ZIKV viral load in urine (median of copies/mL) | 2079 | 2049 | NA | NA |
| NS1/IgM DENV detection | 0 (0.0) | 0 (0.0) | 3 (21.4) | 0 (0.0) |
| DENV qRT-PCR | 0 (0.0) | 0 (0.0) | 0 (0.0) | 0 (0.0) |
| CHIKV qRT-PCR | 0 (0.0) | 0 (0.0) | 0 (0.0) | 0 (0.0) |
| OROV qRT-PCR | 0 (0.0) | 0 (0.0) | 0 (0.0) | 0 (0.0) |
| **Neurological diagnosis** |  |  |  |  |
| Guillain-Barré syndrome | NA | 5 (26.3) | 3 (21.4) | NA |
| Encephalitis | NA | 6 (26.3) | 7 (50) | NA |
| Meningitis | NA | 4 (31.6) | 2 (14.2) | NA |
| Meningoencephalitis | NA | 3 (15.8) | 2 (14.2) | NA |
| Transverse myelitis | NA | 1 (2.6) | 0 (0) | NA |
| Time from onset of viral symptoms to neurological symptoms, median (range) in days | NA | 4 (2 – 15) | 7 (1-29) |  |

### Increased Gas6 expression in ZIKV adult patients with neurological complications

To correlate Gas6 levels to the pathogenesis of ZIKV-associated neurological complications, we first determined Gas6 levels in patients’ serum by ELISA. Figure 1A shows that circulating levels of Gas6 are significantly increased in ZIKV-infected patients compared to healthy donors (Non-Neuro^ZIKV^: 22.56 ng/mL [25-75 interquatile range (IQ) 14.6-29.9] versus 13.36 ng/mL [25-75 IQ 11.6-16.1] in HDs, p = 0.0062), whose circulating Gas6 levels were comparable to previous studies [19–21]. Importantly, Neuro^ZIKV^ patients showed a further increase of serum Gas6 in comparison to Non-Neuro^ZIKV^ patients (33.05ng/mL [25-75IQ 22.5- 40.2], p = 0.0289) (Figure 1A). To rule out the possibility of co-infections influencing Gas6 levels observed in these patients, we used a high-throughput screening (HTS) virus metagenomic approach to identify viral co-infections that were not detected by RT-qPCR or diagnosed by laboratory tests. ZIKV mono-infections in the Neuro^ZIKV^ patients were confirmed in 9 out of 10 patients tested. In one patient, the complete genome of a strain of Pegivirus C (*Pegivirus* genus, *Flaviviridae* family) was sequenced (Supp Fig 1). Noteworthy, this virus has not been associated with neurological complications in humans. Finally, unchanged Gas6 levels in the serum of 14 patients with neurological diseases non-related to acute ZIKV infection (Neuro^NON-ZIKV^) revealed that Gas6 upregulation is a ZIKV-specific response (Figure 1A). In addition, genotyping analysis revealed no difference in *GAS6* haplotypes between the ZIKV-infected patient groups, both showing a predominant frequency of the c.834 + 7G>A AA genotype (Figure 1B).

**Figure 1:**
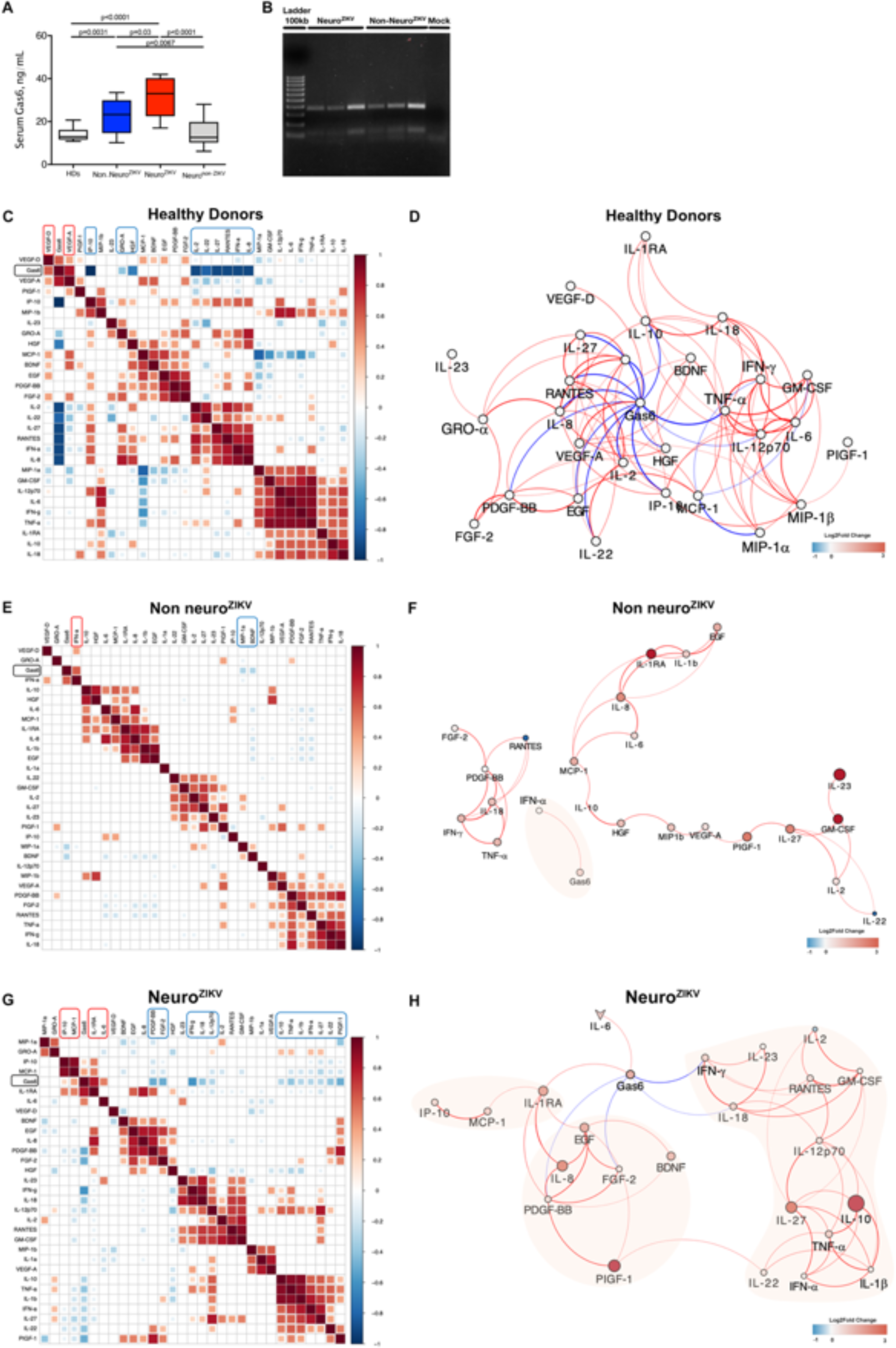
Elevate Gas6 levels in ZIKV patients is a ZIKV-specific response associated with immune signatures related to disease manifestation. (A) Levels of Gas6 in acute-phase patient serum samples of Zika virus (ZIKV) cross-sectional cohort in Campinas, Brazil, were determined by ELISA: Healthy donors (HDs, n =13), ZIKV patients without neurological complications (Non-Neuro^ZIKV^; n = 57), patients with neurological complications (Neuro^ZIKV^; n = 19) and patients presenting neurological complications unrelated to ZIKV infection (Neuro^NON-ZIKV^; n = 14). Gas6 concentration is depicted as Tukey box plots. *P* values were determined using Kruskal-Wallis test with post hoc correction for multiple testing using the original FDR method of Benjamini and Hochberg. (B) Identification of *GAS6* 834 + 7AA genotypes in ZIKV-infected patient groups by agarose gel electrophoresis. Lane 1: Ladder 100kb size marker. Lanes 2-4: Neuro^ZIKV^; Lanes 5-7: Non-Neuro^ZIKV^; Lane 8: Mock. AA homotype showing 345 and 136 bp bands. (C, E and G) Representative images of Pearson’s correlation matrix calculated for each clinical group. A reduced complexity model was established by focusing on significant interactions between ZIKV-specific immune signatures and Gas6 determined by Pearson’s correlation coefficients. Only correlations with associated *p*-value <0.05 are shown and hierarchical clustering was applied. (D, F and H). Representative images of the networks of interactions (prefuse-force layout) determined by Pearson’s correlation coefficients. Each connecting line (edge) represents a significant interaction (*p* < 0.05 and absolute Pearson’s r ≥ 0.5) calculated by the network analysis using the R software. Edge weights and colour are defined as the correlation strength; positive correlations are represented by red edges; negatives correlations are represented by blue edges. Node colour and size represent Log_2_ Fold Change of each biomarker normalized by baseline levels in healthy donors.

### Gas6 upregulation suppresses antiviral IFN response in ZIKV adult patients with neurological complications

To determine the mechanisms by which ZIKV-induced Gas6 upregulation contributes to the pathogenesis of neurological complications in adult patients, we used network analysis of biomarkers and transcriptional profiling to determine how Gas6 orchestrates with a specific signature of immune mediators associated with ZIKV infection, previously determined in our cohort [9]. In accordance with its role as pleiotropic inhibitor of innate immune responses [12, 13, 16], Gas6 negatively correlates with several pro-inflammatory cytokines/chemokines, such as IL-2, IL-8, IL-27, RANTES, IP-10 and TNF-α (r > 0.7; p < 0.05) in healthy donors (Figure 1C, D). Interestingly, a striking change in the pattern of interactions between all biomarkers appeared in Non-Neuro^ZIKV^ patients. In these patients, Gas6 significantly and positively correlates only with IFN-α (r = 0.60; p < 0.05), apart from other 2 functional clusters of significant interactions between the measured biomarkers (Figures 1E, F). In addition, the network graph of Non-Neuro^ZIKV^ patients is more heterogeneous and shows a decentralized topology with lower complexity and connectivity between the immune mediators when compared to the highly dense, homogenous and centralized graph of HDs (Non-Neuro^ZIKV^ vs HDs network density: 0.120 vs 0.224; network centralization: 0.107 vs 0.335; network diameter: 10 vs 5) (Figure 1D, F). Interestingly, a further increase on Gas6 levels above certain threshold (estimated to be above 30ng/mL), as observed in Neuro^ZIKV^ patients (Supp Fig 2), also changes the patterns of interactions. In these patients, Gas6 displays positive correlation with IL-1RA, IL-6, MCP-1 and IP-10 (r = 0.8; 0.5, 0.4 and 0.3, respectively; p<0.05) and negative correlations with IFN-α, IFN-γ and IL-18 (r = -0.4; -0.7 and -0.5, respectively; p<0.05) (Figures 1G, H). In addition, as shown in the network graph (Figure 1H), because IFN-γ forms a functional cluster of interactions with a variety of cytokines with key roles in the antiviral response, Gas6 also displays indirect negative correlations with TNF-α, IL-1β, IL-12, IL-22 and IL-27 (r between -0.3 and -0.5; p < 0.05), as shown in Figure 1G.

**Figure 2:**
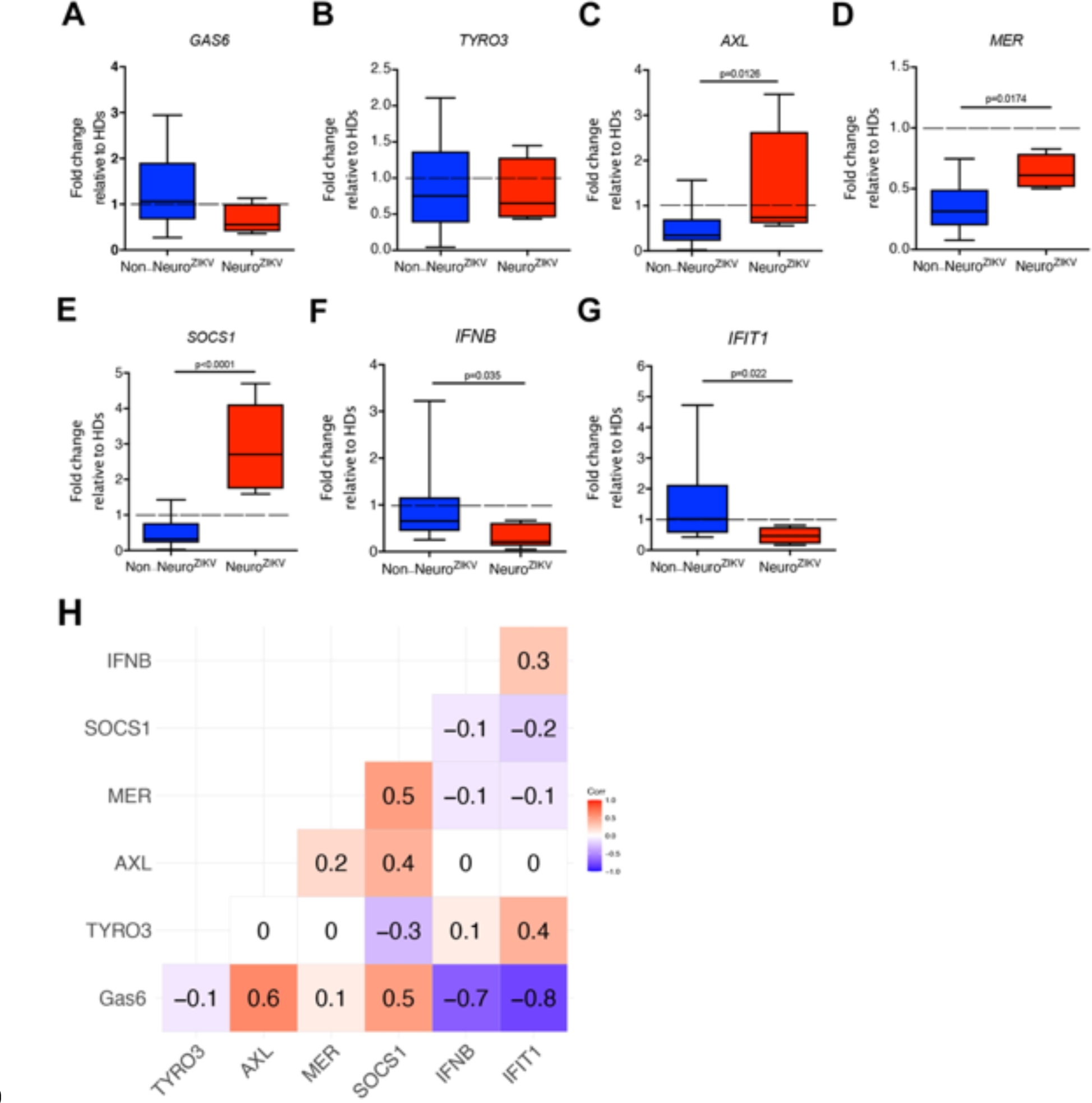
Transcriptional landscape of peripheral blood cells from ZIKV adult patients. Quantitative mRNA expression was determined in peripheral blood cells isolated from healthy donors (HDs) and ZIKV-infected adult patients with (Neuro^ZIKV^) and without neurological complications (Non-Neuro^ZIKV^) by qRT- PCR. (A) *GAS6*; (B) *TYRO*; (C) *AXL*; (D) *MER*; (E) *SOCS1*; (F) *IFNB*; (G) *IFIT1*. Graphs depict fold change calculated relative to healthy donor samples (dashed line) as Tukey box plots, after results were normalized to GAPDH housekeeping gene expression. *P* values were determined using student’s t-test was used to compare groups with normally distributed data or Mann-Whitney test to compare groups with non-normal distributions; \**p* < 0.05; \*\**p* < 0.01. (H) Spearman’s Rank Correlations were determined to assess the association between levels of Gas6 in the serum and expression levels of the transcripts quantified in the matched peripheral blood cells form ZIKV-infected adult patients.

To confirm whether Gas6 drives suppression of antiviral response, we conducted quantitative mRNA expression analysis in peripheral blood cells isolated from healthy donors and patients with and without neurological complications. As shown in Figure 2, although Gas6 mRNA expression did not change (Figure 2A), mRNA expression of Axl and Mer is upregulated in the peripheral blood cells from Neuro^ZIKV^ patients (Figure 2C, D), whereas Tyro3 did not change (Figure 2B). In accordance with TAM-induced SOCS upregulation [13, 22], we observed a significant increase of SOCS1 mRNA in circulating cells from Neuro^ZIKV^ patients (Figure 2E). Conversely, mRNA expression of IFN-β and IFIT-1 is significantly reduced in Neuro^ZIKV^ patients (Figure 2F, G). Thus, in ZIKV adult patients, increased circulating Gas6 levels shows a negative correlation with *IFNB* and *IFIT1* gene expression and positively correlates with *AXL* and *SOCS1* expression in peripheral blood cells (Figure 2H).

### Gas6-producing monocytes dampen the antiviral response during ZIKV infection

Next, we sought to investigate the peripheral cellular sources of Gas6 contributing to its increased levels in the circulation during infection. For that, we cultured and infected PBMCs isolated from healthy donors with ZIKV (Brazilian strain BeH823339) at multiplicities of infection (MOI) 0.1, 1 and 10. Cell pellets and their respective supernatants were collected at different time points post-infection and Gas6 levels were determined by ELISA and cell pellets were used for mRNA quantification by qRT-PCR. PBMCs show an interesting ZIKV viral load-dependent increase of Gas6 production in the supernatants, both at 24 and 48 h post- infection (Figures 3A-C). As neurological complications of ZIKV infection may correlate with BBB disruption, we also evaluated (hBMEC). As shown in Supp Fig 3B, ZIKV infection does not stimulate Gas6 production by hBMECs, indicating that blood mononuclear cells might be the main source of circulating Gas6 during ZIKV infection.

**Figure 3:**
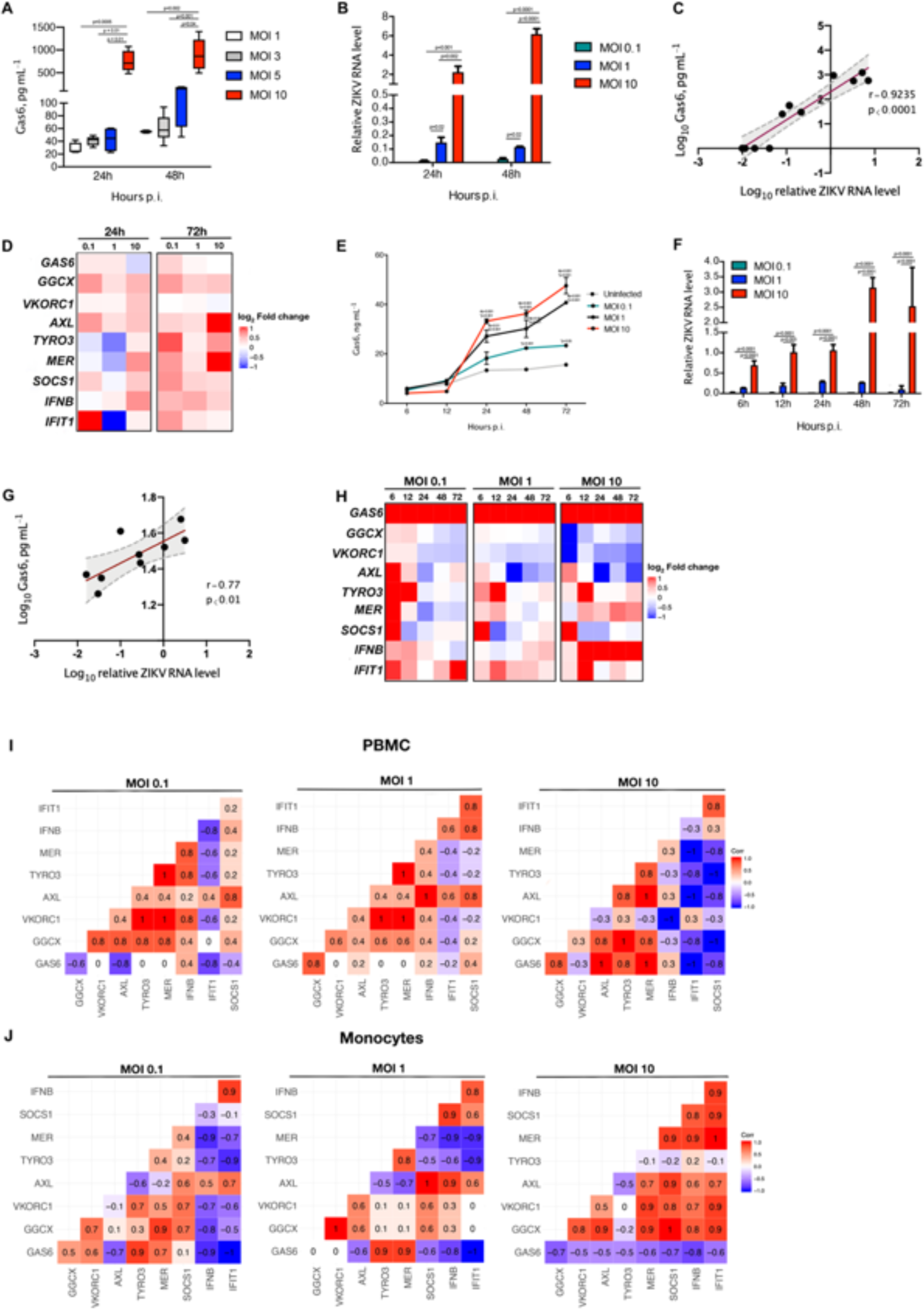
ZIKV Infection stimulates Gas6 upregulation and leads to downregulation of type I IFN response in peripheral blood mononuclear cells (PBMCs) and monocytes. (A) Gas6 levels were determined by ELISA in the supernatant of PBMCs from healthy donors collected at 24h and 48h after *in vitro* infection with ZIKV atdifferent multiplicities of infections (MOI 0.1, 1, 3, 5 and 10). Gas6 concentration is depicted as Tukey box plots. Gas6 was not detected in uninfected (Mock) and ZIKV-infected PBMCs at MOI 0.1, so these conditions are not represented in the graph. The data shown are representative images of two independent experiments using cells from different donors. P values were determined for comparisons between MOIs at the corresponding time point using Kruskal-Wallis test with post hoc correction for multiple testing using the original FDR method of Benjamini and Hochberg. (B, D, F and H) PBMCs and THP-1 monocytes were challenged with ZIKV (MOI 0.1, 1 and 10), respectively. Total cellular RNA was extracted at different time points after infection as indicated in the graphs, and relative viral RNA levels, *GAS6*, *GGCX*, *VKORC1*, *AXL*, *TYRO3*, *MER*, *SOCS1*, *IFNB* AND *IFIT1* mRNA levels were determined by real-time quantitative PCR. (B, F) The data shown are mean ± SEM representative of three independent experiments using cells from different donors. P values were determined for comparisons between conditions at the corresponding time point using One-way analyses of variance statistical test with Bonferroni-corrected multiple comparisons test. (D, H) Heatmap reflecting expression intensity (log_2_ Fold change), in comparison to mock (uninfected cells), of genes in PBMCs or in THP-1 monocytes at different time points (from 6 up to 72h) after *in vitro* infection with ZIKV at different multiplicities of infections (MOI 0.1, 1 and 10). (E) Gas6 levels were determined by ELISA in the supernatant of THP-1 monocytes at different time points (from 6 up to 72h) after *in vitro* infection with ZIKV at different multiplicities of infections (MOI 0.1, 1 and 10). Gas6 concentration is depicted as mean ± SEM. The data shown are representative of three independent experiments. P values were determined using Two-way analyses of variance with Bonferroni-corrected multiple comparisons test. (C, G) Correlation plots comparing logarithmically transformed ZIKV RNA levels with Gas6 concentration within matched cell and supernatant from PBMCs or THP-1 monocytes, respectively, after *in vitro* infection with ZIKV. Regression line indicated in red with 95% confidence area shown in shaded gray. (C) Spearman’s Rank and (G) Pearson’s correlation coefficient and associated p-values are shown. (I, J) Representative images of Spearman’s Rank Correlations matrixes. Correlations were determined to assess the association between transcripts quantified in the PBMCs or in THP-1 monocytes, respectively, after *in vitro* infection with ZIKV at different multiplicities of infections (MOI 0.1, 1 and 10). Relative gene expression levels at different time points (from 6 up to 72h) in the corresponding MOI condition were used as input.

To determine whether increased Gas6 production correlates with transcriptional changes induced by infection, we first verified that Gas6 mRNA is not significantly increased at 24h and 48h p.i., as represented in the heatmap of log2 Fold change in Figure 3D. This is consistent with the analysis ex vivo, as shown in Figure 2A. As γ -glutamyl carboxylation of Gas6 via vitamin-K-dependent enzyme γ-glutamyl carboxylase (GGCX) and Vitamin-K epoxide reductase enzyme complex 1 (VKORC1) [23] is needed for Gas6 biological activity and to bridge enveloped viruses to TAM receptors [23–27], we checked the expression of related molecules. Upregulation of GGCX mRNA expression is observed 24h and 72h p.i in infected cells at all MOIs, while VKORC1 mRNA expression was not affected by infection (Figure 3D, Supp Fig 4A). In agreement with a previous study [28], ZIKV infection induces upregulation of Axl, Tyro and Mer transcripts at higher MOI 10 (Figure 3D, Supp Fig 4A). SOCS-1 is upregulated in ZIKV-infected PBMCs at all MOIs (Figure 3D, Supp Fig 4A), except for MOI 1 at 24 h. Interestingly, although IFN-β mRNA is upregulated later during infection (72h p.i.), the cells seem irresponsive, once IFIT-1 mRNA expression does not change in comparison to uninfected cells at MOIs 1 and 10 (Figure 3D, Supp Fig 4A). Only at the lower viral load (MOI 0.1) at 24h p.i. a significant upregulation of IFIT-1 mRNA expression is observed (Figure 3D, Supp Fig 4A), which later decreases at 72h.

**Figure 4:**
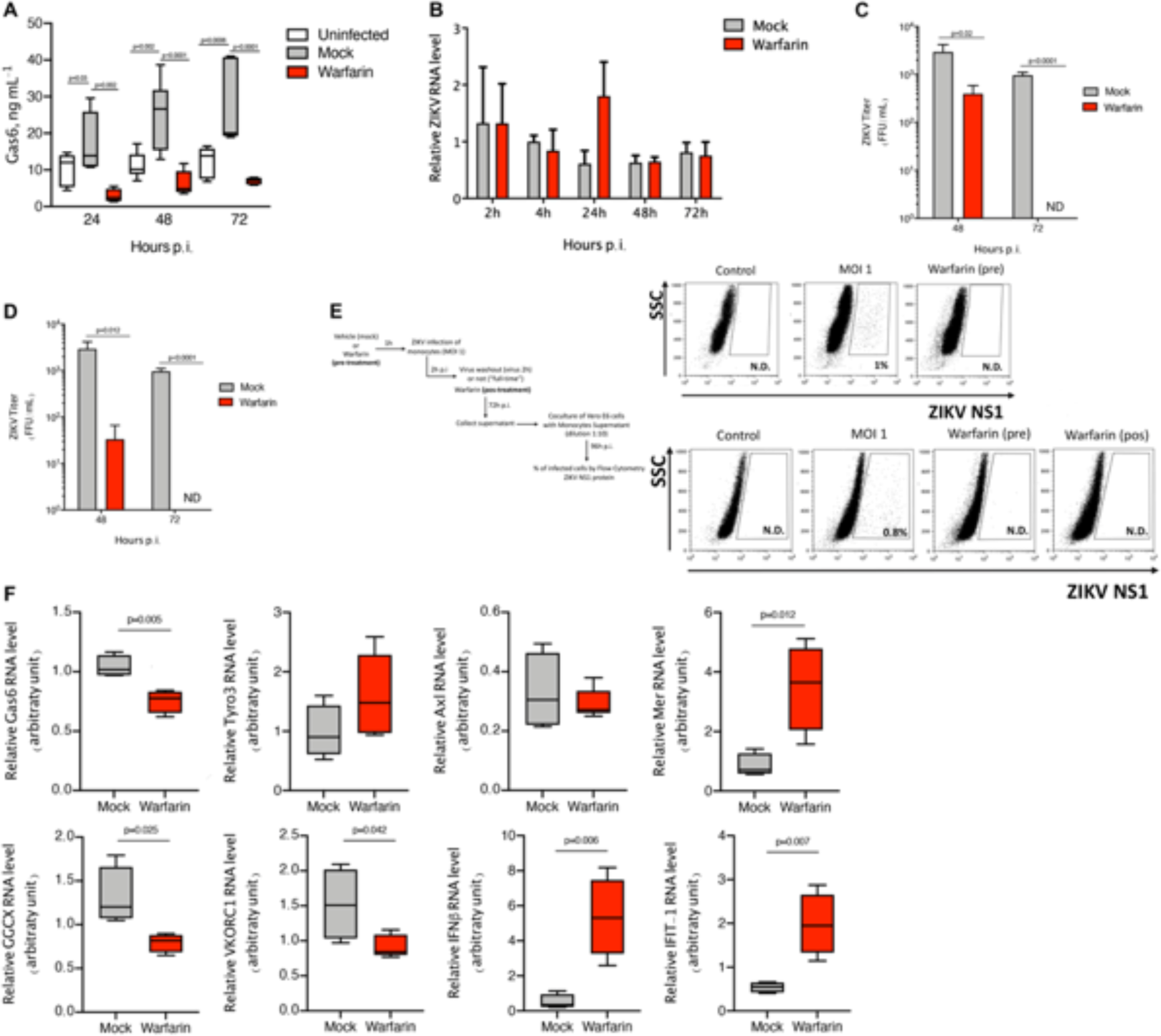
Inhibition of Gas6 γ-glutamic acid carboxylation (Gla domain) by warfarin treatment restores antiviral response. (A) Gas6 levels were determined by ELISA in the supernatant of THP-1 monocytes collected at 24h, 48h and 72h after *in vitro* infection with ZIKV (MOI 1). Infected cells were treated or not (mock) with 2µM warfarin throughout the course of infection. Gas6 concentration is depicted as Tukey box plots. The data shown are representative images of three independent experiments. P values were determined for comparisons between conditions at the corresponding time point using One-way analyses of variance statistical test with Bonferroni-corrected multiple comparisons test. (B, F) THP-1 monocytes were challenged *in vitro* with ZIKV (MOI 1) and treated or not (mock) with 2µM warfarin throughout the course of infection. Total cellular RNA was extracted at different time points after infection. Relative viral RNA levels, *GAS6*, *GGCX*, *VKORC1*, *AXL*, *TYRO3*, *MER*, *SOCS1*, *IFNB* AND *IFIT1* mRNA levels were determined by real-time quantitative PCR. The data shown are representative images of three independent experiments. (F) Graphs depict relative RNA expression as Tukey box plots after results were normalized to GAPDH housekeeping gene expression. P values were determined using student’s t-test to compare groups with normally distributed data or Mann-Whitney test to compare groups with non-normal distributions. (C, D) ZIKV titer (FFU/mL) was determined by focus-forming assay after incubation of ZIKV-permissive Vero E6 cell line with the 48h or 72h supernatant from monocytes after infection *in vitro* with ZIKV (MOI 1), treated or not (mock) with 2µM warfarin at the moment of infection (pre-treatment) or 2h after infection (post-treatment), respectively. The data shown are representative images of three independent experiments. P values were determined using Mann-Whitney test comparing conditions at the corresponding time point. (E) Vero E6 cells were incubated for 4 days with 72h p.i. supernatants from THP-1 monocytes infected *in vitro* with ZIKV MOI 1, treated or not (mock) with 2µM warfarin 1h before infection (pre- treatment) or 2h after infection (pos-treatment). The virus was left throughout the course of the experiment, up to 72h in the monocyte culture (upper panel), or removed 2h after infection by washing the cells three times with PBS (virus washout; lower panel). The percentage of infected cells was determined by flow cytometry analysis of ZIKV NS1 protein expressed in Vero cells. The data shown are representative images of three independent experiments.

Monocytes are the dominant cell type infected by ZIKV among peripheral blood cells [29, 30]. To further understand the role of Gas6 in facilitating ZIKV replication, we used cultured human monocytes (THP-1 cells) infected with ZIKV at MOI 0.1, 1 and 10. Cell pellets and supernatants were collected at different time points after infection. Similarly to PBMCs, Gas6 production by THP-1 cells is increased in a time-dependent and viral load-dependent manner, with a significant increase observed after 24hp.i. (Figure 3E-G). Interestingly, as shown in the heatmap of log2 Fold change in comparison to mock cells (uninfected) in the Figure 3H, Gas6 mRNA is upregulated in THP-1 cells as soon as (6h p.i.) and remains increased throughout the experiment (Supp Fig 4B). GGCX and VKORC1 mRNA expression were not significantly altered, although it tends to decrease at higher MOI 10 (Figure 3H, Supp Fig 4B). Importantly, Axl, Mer, Tyro3 and SOCS-1 mRNA expression was increased early after infection (6-12h p.i.), in particular at lower MOI 0.1 (Figure 3H, Supp Fig 4B). IFN-β is upregulated only at MOI 10 (Figure 3H, Supp Fig 4B), but the cells are irresponsive, once IFIT-1expression does not significantly change in comparison to uninfected cells (Figure 3H). Similar to PBMCs, a late significant increase (72h p.i.) of IFIT-1 mRNA expression is observed only at the lower MOI 0.1 (Figure 3H, Supp Fig 4B). Interestingly, in PBMCs, Gas6 expression negatively correlates with IFIT-1 at all viral loads (Figure 3I) while in monocytes THP-1 cells, Gas6 negatively correlates with IFN-β and IFIT-1 at all viral loads (Figure 3J). Altogether, these data show that ZIKV infection induces a significant upregulation of Gas6 production in monocytes, which in turn facilitates viral replication by suppressing type I IFN antiviral response.

### Inhibition of Gas6 γ-glutamic acid carboxylation (Gla domain) restores antiviral response

To further determine the mechanistic link between Gas6 production, ZIKV replication and suppression of antiviral response, we tested whether inhibiting Vitamin-K-dependent γ- carboxylation of Gas6 glutamic acid residues (Gla domain) restores antiviral immune response and controls viral replication. We used low-dose warfarin, which specifically block γ- carboxylation of Gla domain in Gas6, resulting in decreased TAM receptor activation [23, 27, 31]. In parallel, we used R428, a well-known inhibitor of Axl tyrosine kinase activity tested in ZIKV infection [16]. THP-1 cells were infected with ZIKV at MOI 1 and treated with 2μM of warfarin or 2μM R428 throughout the course of infection (up to 72h). Treatments started at the moment of infection or 2h p.i. (after virus entry), and cell pellets and their respective supernatants were collected at different time points. Warfarin treatment blocked Gas6 upregulation at the protein (Figure 4A) and mRNA levels (Figure 4F), although ZIKV RNA level did not change (Figure 4B). Warfarin also decreased GGCX and VKORC1 mRNA expression (Figure 4F). Corroborating this, R428 treatment potently reduced ZIKV-induced Gas6 production (Supp Fig 5A).

**Figure 5:**
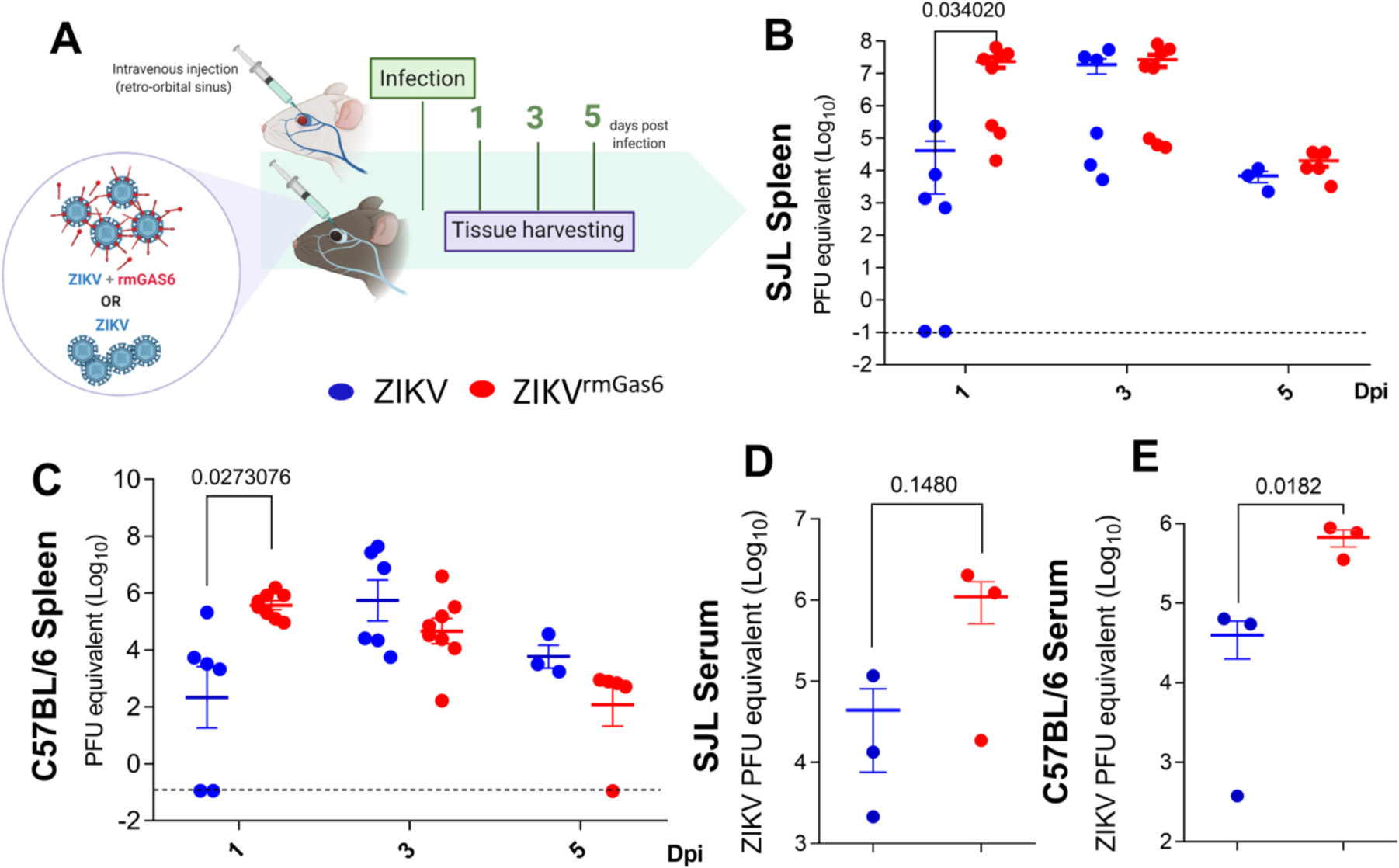
Gas6 facilitates ZIKV replication in immunocompetent mice. SJL and C57BL/6 mice were infected by intravenous route with 10^2^ pfu of pure ZIKV (BeH815744) or ZIKV previously incubated with rmGas6 (1µg/mL) (ZIKV^rmGas6^) (A). After 1-, 3- and 5-days post-infection the viral load was analysed by qPCR in spleen (B, C) and serum (D, E) of both mice strains. *P* values were determined using a Student’s *t*-test. Graph bars are shown as mean ± SEM and are representative of three independent experiments; 1 and 3 dpi: ZIKV (n = 6), ZIKV^rmGas6^ (n = 8); 5 dpi: ZIKV (n = 3), ZIKV^rmGas6^ (n = 5).

Interestingly, incubation of ZIKV -permissive Vero cell line with the supernatant from the ZIKV-infected monocytes revealed that warfarin treatment, at the moment of infection or 2h p.i., significantly decreases the production of viable viral particles at 48h p.i. (Figures 4C, D). At 72h p.i., the drug completely blocked viral production (Figures 4C, D). R428 did not change ZIKV RNA levels but showed a modest but significant effect on decreasing viral production at 72h p.i. (Supp Fig 5B, C). Accordingly, flow cytometry analysis of ZIKV NS1 protein in Vero cells showed that infection was not detected when cells were exposed to supernatants from monocytes treated with warfarin, regardless whether the initial virus inoculum was washed-out 2h p.i. or not or whether warfarin treatment started 2h p.i. (pos-treatment approach) or at the moment of infection (pre-treatment approach) (Figure 4E). Interestingly, warfarin induced a potent upregulation of IFNβ and IFIT-1 expression (Figure 4F). Meanwhile, R428 increased the expression of Axl and Mer (Supp Fig 5D). Consistent with the blockade of type I IFN by Axl signalling pathway, R428 allowed the increase of IFN-β and IFIT-1 (Supp Fig 5D), while it did not significantly affect GGCX or VKORC1 mRNA expression (Sup Fig 5D). Thus, blockage of Gas6 activity by inhibition of γ-carboxylation of Gla domain restores type I IFN antiviral response sufficient to block the production of infective virus particles. Collectively, these data show the mechanism by which Gas6 upregulation contributes to a higher ZIKV infection.

### Gas6 facilitates ZIKV replication in immunocompetent mice

So far, our data show that Gas6 expression is stimulated during ZIKV infection and it is associated with neurological complications in human patients. The underlying pathogenic mechanism of Gas6 involves attachment and invasion and facilitation of viral replication by suppressing the antiviral response. Although we prove a correlation between Gas6 levels, ZIKV RNA load and production of infecting particles in infected cells *in vitro*, we could not find the same association in our cross-sectional cohort. To confirm that upregulation of Gas6 favours viral replication *in vivo*, immunocompetent adult C57BL/6 and SJL mice were intravenously infected with 10^2^ pfu ZIKV (BeH815744) previously incubated or not with 1μg/mL recombinant mouse Gas6 (ZIKV^rmGas6^) (Figure 5A). Viral load was determined at 1, 3 and 5 d.p.i. in the spleen and serum (Figure 5A). Interestingly, infection with Gas6-coated ZIKV resulted in increased viral RNA in the spleen (Figure 5B-C) and serum (Figure 5D-E) in both strains at 1 d.p.i. After 3 and 5 days of infection, differences were no longer detected.

Similar to *in vitro* infection of cultured human cells, these data corroborate the role of Gas6 in favouring ZIKV infection and replication.

### Gas6 promotes congenital syndrome and transplacental infection in ZIKV-infected immunocompetent mice

To further evaluate whether Gas6 drives disease severity during ZIKV infection, we evaluated its association with transplacental infection and development of congenital zika syndrome (CZS). Determination of Gas6 levels in the serum of 8 ZIKV-infected pregnant women revealed a significant increase during acute phase of infection (18.05 ± 1.97 ng/mL versus 13.36 ± 1.23 ng/mL) (Supp Figure 6A). On the other hand, we did not detect a difference in Gas6 levels in the serum of babies with (n = 2) or without (n = 4) CZS (Supp Figure 6B).

**Figure 6:**
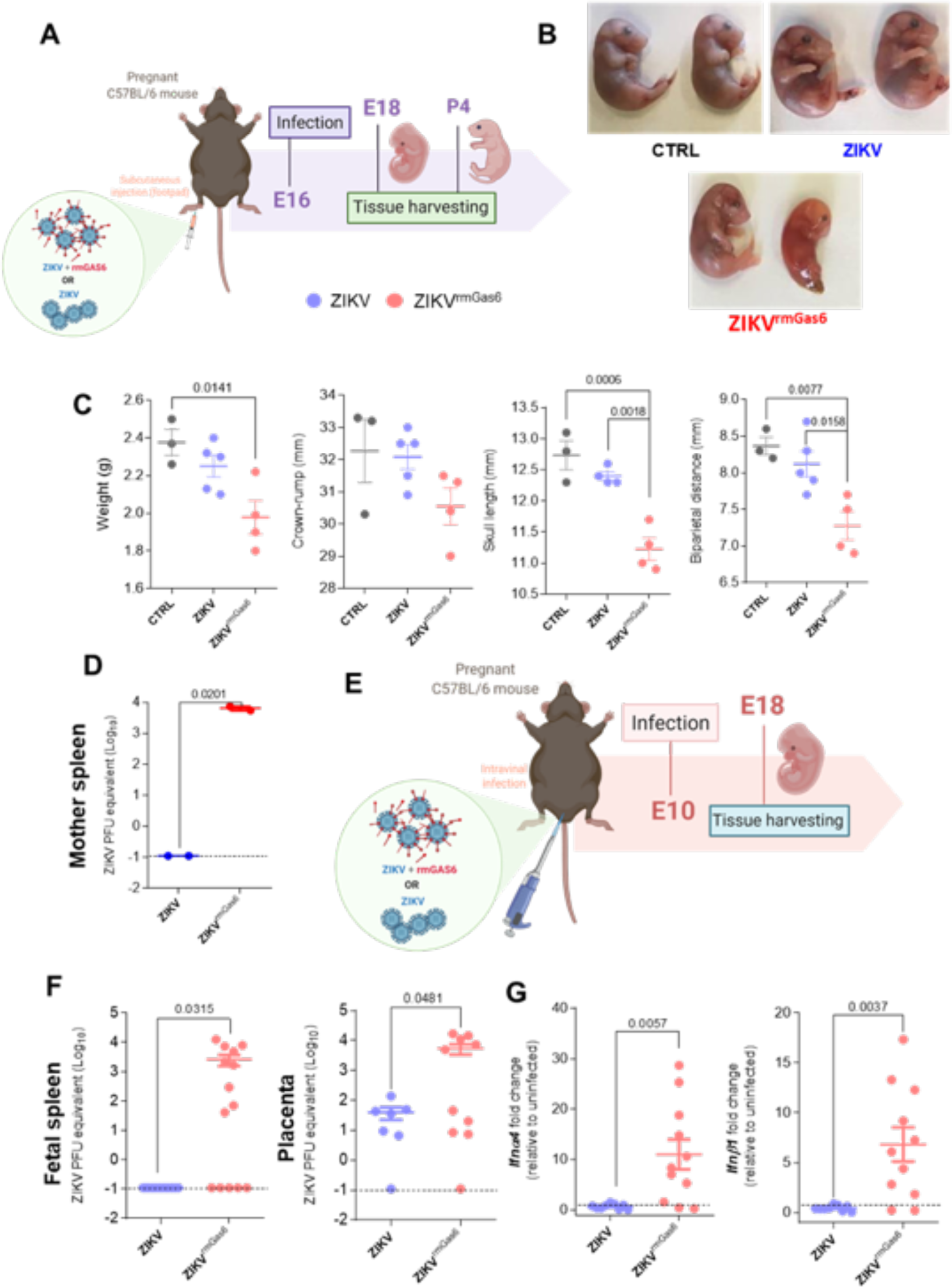
Gas6 promotes ZCS and transplacental infection in ZIKV-infected resistant immunocompetent mice. Pregnant mice were infected subcutaneously with pure ZIKV (10^5^) or ZIKV previously incubated with rmGas6 (1µg/mL) (ZIKV^rmGas6^) on E (embryonic day) 16. The organs were harvested on E18 or postpartum day (P) 4 (A). The viral load was analysed by qPCR. Foetus’s pictures at E18 (B). Dimension analyses at P4 (C). *P* values were determined using a one-way ANOVA followed by Tukey’s post hoc test. Mother’s spleen viral load (D). Pregnant mice were also infected intravaginally on E10 with pure ZIKV (10^5^) or ZIKV previously incubated with rmGas6 (1µg/mL). Tissues were harvested at E18 (E). Viral load in spleen and placenta (F). Gene expression in placenta (G). Fold change was calculated between uninfected and infected groups. The significances between the groups were performed by Student *t*-test (D, F and G). Graph bars are shown as mean ± SEM and are representative of two independent experiments. Numbers of experimental groups: (B) Control n=6; ZIKV n=11; ZIKV^rmGas 6^ n=14. (C) Control n=3; ZIKV n=5; ZIKV^rmGas6^ n=4. (D) ZIKV n=2; ZIKV^rmGas6^ n=2. (F) ZIKV n=7; ZIKV^rmGas6^ n=10.

As the number of ZIKV-infected pregnant women with fetal growth-associated malformations included in this study was limited to two patients, we decided to evaluate the effect of Gas6 using a mouse model of CZS. For this, we subcutaneously injected pregnant C57BL/6 mice with Gas6-coated ZIKV (ZIKV^rmGas6^) at embryonic day (E) 16 and performed analysis at E18 or postnatal day (P) 4 (Figure 6A and Supp Figure 6). Importantly, infection of pregnant C57BL/6 mice with ZIKV^rmGas6^ rendered their offspring susceptible to fetal malformations (Figure 6B). We evidenced macroscopic changes at E18 and growth delay at P4 (Figure 6B, C). Significant differences in biparietal distance, skull length and weight of the ZIKV^rmGas6^ newborns were observed (Figure 6C). No difference in viral load between ZIKV and ZIKV^rmGas6^ groups was found in the placenta and fetal spleen at E18 (Supp Figure 7A) as well as in the spleen of P4 newborns (Supp Figure 7B). However, a higher viral load was observed in the spleen of ZIKV^rmGas6^-infected pregnant mice as long as 8 days post-infection (day 0 at E16, and day 8 at P4) (Figure 6D).

**Figure 7:**
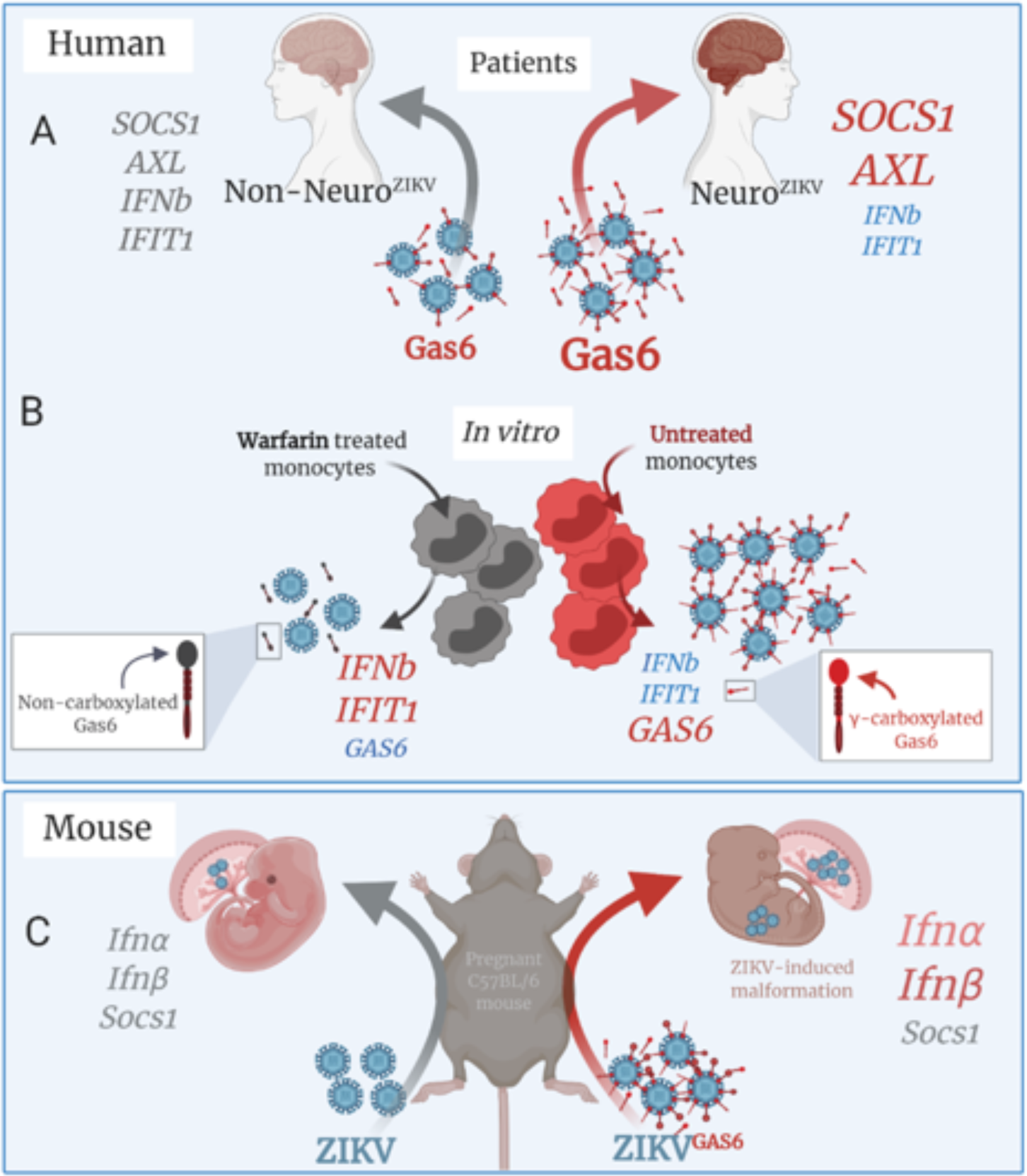
Gas6 is associated with neurological complications in humans and drives ZIKV- congenital malformations in immunocompetent mice. (A) Elevated Gas6 levels in the serum of ZIKV patients with neurological complications (Neuro ZIKV) is associated with upregulation of AXL and SOCS1 and downregulation of IFNβ and IFIT1. (B) In vitro treatment with Warfarin reduces Gas6 levels and restores type I IFN antiviral response by inhibiting Gas6 γ-glutamic acid carboxylation thus decrease ZIKV infection. (C) The mouse infection with ZIKV pre-treated with recombinant Gas6 (ZIKV Gas6) facilitates Zika virus infection and leads to malformations in the offspring which is associated with upregulation of Ifnα and Ifnβ. Created with BioRender.com.

Since Axl is expressed in decidua, Hoffbauer, trophoblast and fibroblasts cells of human placenta^2^ and ZIKV has been considered a sexually transmitted disease (STD), we sought to verify whether Gas6 facilitates intra-uterine ZIKV infection. For that, C57BL/6 pregnant mice were intravaginally infected at E10 with ZIKV or ZIKV^rmGas6^ and viral load was determined at E18 (Figure 6E). Contrasting with the subcutaneous infection, no difference in fetal size was observed (Supp Figure 7C, D), but there is a significant increase in ZIKV load in the fetal spleen and in the placenta of ZIKV^rmGas6^-infected pregnant mice (Figure 6F). Although we see no change in *Stat1* and *Socs1* mRNA expression, *Ifnα4 and Ifnβ1* expression were unexpectedly increased in the placenta of ZIKV^rmGas6^-infected pregnant mice (Figure 6G).

## DISCUSSION

The ZIKV epidemic that occurred in 2016 was a great public health problem. ZIKV is able to cross the blood-brain and placental barriers, infecting the central nervous system (CNS) of adults and developing foetuses, greatly increasing the severity of the disease and the impact of this infection [32–39]. In fact, studies with *ex vivo* slices from human cortical tissue and adult mice demonstrated that ZIKV is able to infect and replicate in mature neurons, causing local inflammation and damage in hippocampal synapses, resulting in impairment of synaptic function and memory [40]. These findings show that contemporary occurring ZIKV strains are highly neurotropic and detrimental to both mature as well as to the developing central nervous system (CNS) [33, 41]. Although studies have raised the contribution of several factors in the development of these conditions [34, 42, 43], the mechanisms by which the virus reaches the brain of adult individuals, is not fully understood. In this context, here we demonstrate that circulating levels of Gas6 are upregulated in ZIKV-infected patients showing neurological complications. This upregulation is ZIKV-specific and could be influenced by intrinsic host features such as allelic variability of the human *GAS6* gene. Elucidation of the human *GAS6* gene structure and allelic variants revealed the presence of eight different variants confirmed to be single nucleotide polymorphisms (SNPs). The SNP in the intron 8 (c.834+7G>A; genotypes GG, AG, and AA) controls circulating levels of Gas6 and plays a key role in the pathogenesis of different circulatory disease, such as preeclampsia, acute coronary syndrome and stroke [21, 44–46]. We observed no difference in *GAS6* haplotypes, as both groups of patients showed the predominant frequency of the c.834 + 7G>A;AA genotype, indicating that higher Gas6 levels observed in Neuro^ZIKV^ patients correlate to ZIKV infection.

Different from hBMECs, monocytes were more responsive to ZIKV infection by upregulating transcriptional and translational machinery resulting in the increase of Gas6 expression. This indicates that monocytes might be the main cellular source and contributing to increased Gas6 levels in the plasma of ZIKV-infected patients. It is worth mentioning that this is consistent with the fact that circulating CD14^+^CD16^+^ monocytes carry ZIKV particles during infection [29, 30]. Moreover, in immune cells, engagement of TAM receptors by Gas6 results in the inhibition of inflammatory responses driven by Toll-like receptors (TLR) and type I IFN signalling pathways [12-14, 22, 47]. The mechanism depends on the expression of SOCS, resulting in an autocrine and paracrine inhibition of both signalling pathways during ZIKVinfection *in vivo.* This response in leukocytes, in particularly in monocytes, and microvascular endothelial cells might contribute to virus spread and replication in the periphery and potentially in target tissues, such as the CNS. Therefore, our network analysis of associations between Gas6 and ZIKV-specific immune signatures and transcriptional profiling of peripheral blood cells further elucidate the pathogenic mechanism of action of Gas6. Different from healthy donors, the increase of Gas6 levels in Non-Neuro^ZIKV^ patients changes its patterns of correlations, and we have found a significant positive correlation between Gas6 and IFN-α. This along with increased circulating levels of proinflammatory cytokines, chemokines and growth factors suggests that an active production of a network of immune mediators could provide a strong antiviral environment during the acute phase of disease, resulting in a milder clinical outcome. Conversely, in Neuro^ZIKV^ patients, the further increase of Gas6 expression changes its pattern of interactions. Gas6 displays a positive correlation with IL-6, MCP-1, IP-10, IL-1RA and negative correlations with IFN-α, IFN-γ and IL-18. In addition, as shown in the network graph, as IFN-γ forms a functional cluster with a variety of important mediators in host defense against infection [9, 48], Gas6 also displays indirect negative correlations with TNF-α, IL-1β, IL-12, IL-22 and IL-27. This pattern of interactions reveals that elevation of circulating Gas6 levels above a certain threshold, which we estimate to be higher than 30ng/mL, dampens the protective immune response that provide a strong antiviral environment and a milder clinical outcome. In support, transcriptional analysis in peripheral blood cells from ZIKV patients with neurological complications indicate that this response is achieved by a ZIKV–Gas6–TAM receptor interaction that ultimately induces downregulation of type I IFN genes modulated by SOCS1. This mechanism was corroborated in cultured monocytes, where ZIKV-induced Gas6 expression also correlated with suppression of type I IFN response.

The antiviral response also involves recruitment and coordination of specific subsets of immune cells orchestrated primarily by chemokines. Gas6 can also function as an inflammatory molecule by inducing leukocyte adhesion on endothelial cells surface and extravasation through a P-selectin–dependent mechanism [49]. Moreover, Axl and Mer, in cooperation with IFNAR signalling, have been described as key molecules for maintenance of the blood-brain barrier (BBB) and protection to WNV and La Crosse virus infection [50]. Accordingly, *in vivo* studies of ZIKV infection in immunocompromised IFNAR-KO mice lacking Axl have shown protection from ZIKV neuropathogenesis and severe infection [18].

In this context, suppression of IFNAR signalling due to increased amounts of circulating Gas6 in Neuro^ZIKV^ patients potentially impairs BBB homeostasis and integrity. This may allow viral and cellular extravasation to CNS though loose endothelial cells junctions. Noteworthy, in Neuro^ZIKV^ patients, Gas6 positively correlates with IL-6, IP-10 and MCP-1, suggesting that there might be a parallel between ZIKV and cytokine release syndrome (CRS). In addition, our data reveal a unique and inappropriate inflammatory response in Neuro^ZIKV^ patients. This response is defined by a failed type IFN-I response in the periphery, juxtaposed to elevated chemokines levels and high expression of IL-6, leading to recruitment and infiltration of effector immune cells in deep tissues. Thus, infiltration of cells, such as CD4 and CD8 T cells, along with infection of mature neurons could induce a local inflammatory response in the CNS, potentially resulting in neurological complications [40]. We propose that Gas6 mediated reduced innate antiviral defences coupled with exuberant inflammatory cytokine production are driving features of severe clinical outcome in ZIKV adult patients.

It has been shown that inhibition of Axl signalling by different pharmacological or genetic approaches decrease ZIKV replication *in vitro* in CNS cells, endothelial cells and dendritic cells as well as reduce brain pathology in experimental models [16, 18, 51–53]. TAM receptors activation by Gas6 is highly dependent on its γ-carboxyglutamic acid-rich (Gla) domain, required for its biological activity and to bridge enveloped viruses to bind and activate TAM receptors [23–27]. After translation, Gas6 is activated by γ -glutamyl carboxylation via the Vitamin-K cycle and cycle [23]. Vitamin-K epoxide reductase enzyme complex 1 (VKORC1) recycles Vitamin-K Epoxide back to Vitamin-K Hydroquinone, which in turn serves as a co-factor in the γ-carboxylation of Gas6 induced by Vitamin-K-dependent enzyme γ-glutamyl carboxylase (GGCX). Low-dose of warfarin functions as a direct VKORC1 inhibitor, preventing γ-carboxylation of Gas6 and TAM receptor activation [23, 31]. Thus, we used low-dose warfarin to further determine the mechanistic link between Gas6 production, ZIKV replication and suppression of antiviral response. It is important to highlight that one could argue that decreased Gas6 production after warfarin treatment could be a result of decreased binding of anti-Gas6 capture antibody in the ELISA due to restricted recognition of γ-carboxylated residues (amino acids 53-92) in Gas6 protein. However, the capture antibody recognizes the residues 118-678, which are not in the Gla-domain. In addition, Gas6 can be transcriptionally upregulated by Axl-mediated autocrine mechanisms [23], which could explain warfarin-induced downregulation of Gas6 expression. In our experiments, this mechanism was confirmed by restoration of antiviral response and complete blockage of production of infective viral particles when monocytes were treated with warfarin. These findings implicate that, by tethering ZIKV to TAM receptors, Gas6 mediates the pivotal suppression of type I interferon receptor (IFNAR) signalling, thereby favouring ZIKV evasion from antiviral immunity and sustained replication, pointing out how interactions with membrane receptors go beyond attachment and internalization of viral particles [54].

We demonstrate a correlation between Gas6 protein expression, ZIKV RNA load and production of infecting particles associated with suppression of type I IFN response in human cells infected *in vitro*. Although we did not detect difference in ZIKV RNA load in the peripheral blood specimens in our cross-sectional cohort, this could probably be due to the moment blood sampling was performed. When immunocompetent adult C57BL/6 and SJL mice were infected with Gas6-coated ZIKV (ZIKV^rmGas6^), we observed that upregulation of Gas6 increases viral load very early after infection, at 1 d.p.i. After 3 and 5 days of infection, differences were no longer detected. This might explain our observation in patients, as the median interval between illness onset and sampling was 4 days, ranging from 1 to 6 days. Thus, similar to *in vitro* infection of cultured human cells these data corroborate the role of Gas6 in increasing ZIKV replication *in vivo*. Intriguingly, some studies that also used i*n vivo* experimental approaches have demonstrated that the presence of TAM receptors are not required for ZIKV infection in the testis, eye, brain and neither for transplacental infection [55–57]. Nevertheless, our data are the first to describe that Gas6 facilitates ZIKV replication in adult immunocompetent mice model acting as a gatekeeper for viral entry. In addition, these observations corroborate *in vitro* studies of ZIKV and DENV infection [16, 24].

We had previously evidenced the resistance of C57BL/6 mice to congenital malformations caused by ZIKV [34]. Here, we demonstrated that rmGas6 could render C57BL/6 susceptible to ZIKV induced growth restriction and viral transplacental passage associated with increased placenta *Ifnα* and *Ifnβ* expression. Axl is expressed in decidua, Hoffbauer, trophoblast and fibroblasts cells of human placenta, suggesting as the receptor viral transplacental passage [58]. Interestingly, recent studies, one in preprint, have demonstrated that CZS is associated with exacerbated type I IFN and insufficient type III IFN in placenta at term, probably without modulation of TAM or TIM receptor mRNA expression in placental sites infected with ZIKV [59, 60]. In addition, human babies carrying CG/CC genotypes of rs2257167 in IFNAR1 presented higher risk of developing CZS [59]. However, the correlation between TAM receptors and Gas6 expression in susceptibility to congenital ZIKV syndrome has not been established. Our data demonstrate that intravaginal infection with Gas6-coated ZIKV leads to increased viral load in the placenta of pregnant mice and in the spleen of their foetuses, pointing out the role of Gas6-TAM receptors for transplacental passage. Supporting these results, previous studies have demonstrated that placental viral infection is capable to elicit morphological changes in the foetal brain associated with IFN-β immune response at the maternal-foetal interface [58]. Interestingly, deficient *Ifnar* mice, despite higher viral load, do not present significant impairment in the offspring development. However, those whom type I IFN response was present *(Ifnar*^+/-^), despite lower viral load, demonstrated placental spiral arteries apoptosis, foetal hypoxia and prominent growth restriction. In the same study, intraperitoneal injection of poly: IC, a major TLR3 agonist which is also activated by ZIKV [61], led to resorption of all foetuses in an IFNAR-dependent manner [58]. Accordingly, the hypothesis for our findings would be that the immunocompetence described in C57BL/6 animals may be beneficial while low amounts of pathogen are present. However, if viral load is increased, as it happens due to infection with ZIKV^rmGas6^, the exacerbated immune response in the pregnant mother could result in detrimental changes to foetal development probably resulting from increased placental damage.

Together, our observations in ZIKV-infected adult patients and in vitro infections identify a relevant pathogenic mechanism associated to the development of severe outcomes during ZIKV infection, where the virus promotes upregulation of its own ligand Gas6, which contributes to viral infectivity by suppressing an efficient type I IFN antiviral response (Figure 7A and B). Our *in vivo* findings in immunocompetent mice models corroborate the role of Gas6 for favouring ZIKV infection by acting as a gatekeeper in addition to its association with the pathogenesis of congenital syndrome (Figure 7C). Given the novelty of our findings, the relevance to other fields such as the correlation of Gas6 plasma levels and disease severity that until now has been described in lupus [62], liver fibrosis [63] and preeclampsia [64], the relevance of the immunocompetent mouse model for studies with ZIKV and the potential for development of new therapeutic agents that may target Gas6, our data open avenues for research on the factors that drive neurological complications in adults and newborns by this important emerging virus.

## MATERIALS AND METHODS

### Ethical Approval

The present study was conducted according to the Declaration of Helsinki principles after approval by the Research Ethics Committee of the University of Campinas (CAAE: 56793516.0.0000.5404) [9]. Written informed consent was obtained from all participants or from participants’ parents / legal guardians. In addition, animal experiments were carried out in accordance with the recommendations of the IACUC (Institutional Animal Care and Use Committee). The protocols were approved by Ethics Committee for Animal Research of University of Sao Paulo (CEUA - 63/2016) and all efforts were made to minimize animal suffering.

### Patients

This was a cross-sectional study that enrolled 90 patients and 13 healthy donors during the Zika virus (ZIKV) epidemic in Brazil. The patients were admitted to hospitals through emergency departments (ED) in the city of Campinas, Southeast of Brazil, during the ZIKV outbreak (February 2016 to June 2017). 57 of these 90 patients had a mild self-limited illness by ZIKV, characterized by the presence of the following symptoms: fever, rash, conjunctivitis, myalgia, headache, arthralgia and periarticular edema, while 33 patients had clinical signs compatible with neurological syndromes, 19 with diagnosis of ZIKV and 14 without ZIKV detection. This study included patients presenting clinical diagnosis of encephalitis, meningoencephalitis, transverse myelitis and Guillain-Barré syndrome of undetermined origin that had preceding symptoms compatible with arbovirus infection up to 60 days before the onset of the neurological condition. The clinical classification was done following international clinical criteria [65, 66] and patients with history of a prior motor neuropathy or spinal cord disease were excluded of the study. Clinical data were retrospectively retrieved from medical records and the clinical and demographics data are summarized in the Table 1 in the main text. Patients included in this study were grouped as follows:

*Patients ZIKV^+^ with mild symptoms* (Non-Neuro^ZIKV^): 57 patients presenting mild symptoms of infection, such as low fever, rash, myalgia and conjunctivitis. ZIKV infection was confirmed by qRT-PCR.

*Patients ZIKV^+^ with neurological complications* (Neuro^ZIKV^): 19 patients presenting neurologic complications secondary to ZIKV were diagnosed with Guillain-Barré syndrome, encephalitis, meningitis, meningoencephalitis or transverse myelitis according to clinical criteria. Neurological complications started at a median of 4 days after onset of ZIKV acute symptoms. ZIKV infection was confirmed by qRT-PCR and/or specific IgM detection,

*Patients ZIKV^-^ with neurological symptoms* (Neuro^NON-ZIKV^): 14 patients presenting neurologic complications as described above but were negative for ZIKV by qRT-PCR and/or specific IgM detection. Although three of these patients were positive for DENV by NS1/IgM Rapid immunochromatographic tests, the pathological origin of these neurological symptoms was undetermined at the moment of sample collection. All these patients were negative for DENV, CHIKV and OROV by qRT-PCR.

*Healthy donors (HDs):* 13 age-matched individuals without signs of infection within 30 days prior to sample collection. They were included and pre-screened for presence of ZIKV RNA and ZIKV-specific antibodies.

Additionally, not included in the 90 described patients, we analysed acute-phase serum samples of 8 ZIKV-infected pregnant women and 6 infants born of these women. Of these infants, two had CNS abnormalities associated to Congenital Zika Syndrome (CZS), while 4 infants were healthy, without congenital zika syndrome (Non CZS). All participants were tested for a series of arboviruses by qRT-PCR. All patients and healthy donors were negative to Dengue (DENV), Chikungunya viruses (CHKV) and Oropouche were negative as determined by RT-qPCR. Samples were collected after consent of the patients.

### Serum Collection and Processing

Peripheral blood and/or urine specimens were collected at a median of 3 days post-illness onset. Serum was obtained from 10 mL of peripheral blood collected in a dry heparinized tube after peripheral venepuncture. All samples were transported and processed as previously described [9]. Positivity ZIKV RNA in the blood, serum and urine samples was verified by real-time quantitative RT-PCR (qRT-PCR) [9, 67]. In addition, the presence of ZIKV-specific IgM and IgG antibodies in the serum was determined by enzyme-linked immunosorbent assay (ELISA), as previously described [9, 67].

### Cells

This study was conducted using the following cells lines: Vero E6 (ATCC®, (Manassas, Virginia, USA) CRL-1586™), C6/36 (ATCC® CRL-1660™), THP-1 (ATCC® TIB-202™) and hBMECs (previously described here [68] and kindly provided by Dr. Julio Scharfstein – Instituto de Biofísica Carlos Chagas Filho, UFRJ). VeroE6 were cultured in DMEM medium (high glucose) supplemented with 10% fetal bovine serum (FBS). mosquito cell line were cultured in Leibovitz’s L-15 medium supplemented with 10% FBS and 1% penicillin/streptomycin. Human brain microvascular endothelial cells (hBMEC) were cultured in DMEM high glucose, supplemented with 1% L-glutamine, 1% non-essential aminoacids, and 10% FBS [19]. Peripheral blood mononuclear cells (PBMC) from healthy donors were obtained after centrifugation of buffy coat samples over ficoll-hypaque gradient. Subsequently, the mononuclear cell ring was collected and washed with 1x PBS, followed by centrifugation. Then, the pellet was resuspended in complete RPMI-1640 (supplemented with 10% FBS and 1% antibiotics). Cells were then counted for determination of cell viability by Trypan Blue. PBMCs and THP-1 human monocytic cell line were maintained in a complete RPMI-1640 medium supplemented with 10% FBS, 2mM L-Glutamine (Corning), and 1x Penicillin-Streptomycin Solution at 37°C in a fully humidified atmosphere containing 5% CO_2_.

### Virus strains

For *in vitro* studies in cultured human cells, Brazilian ZIKV strain (BeH823339, GenBank KU729217), originally isolated from a patient in Ceará, Brazil in 2015, was provided by Professor Edison Durigon (Biomedical Sciences Institute, University of São Paulo, Brazil). Virus stocks were produced by inoculating Vero CCL81 cells (ATCC) with ZIKV in minimum essential medium (MEM) for 2h at 37°C and 5% CO_2_. Further, the supernatant was removed, and MEM supplemented with 2% FBS, 1% penicillin and streptomycin was added. The cells were incubated for 4 days until 70% of cytopathic effect. Supernatant was then collected, centrifuged for 5 min at 10,000g, 4°C and snap-frozen at -80°C until use.

For *in vivo* studies in mice, Brazilian ZIKV (BeH815744, GenBank KU365780) was provided by Evandro Chagas Institute in Belém, Pará, Brazil and propagated in C6/36 cells. Cultures were infected for 1h at 27ᵒC in the absence of CO_2_. Further, 45mL of complete medium was added (2% of FBS + 1% of Pen/Strep) and cultures were followed until reaching cytopathic effect. At this time, supernatants were harvested and centrifuged at 3200rpm for 10min at 4°C to remove any detached cell. ZIKV culture supernatants were further precipitated with 50% of PEG (polyethylene glycol) for 18h at 4°C. Precipitated virus supernatants were centrifuged (30’, 3200 *g*, 4°C), and the pellet was diluted in DMEM with 25Mm HEPES quantified by PFU assay in VERO cells and used as necessary.

### Mice

SJL and C57BL/6 mice were bred under specific-pathogen-free conditions at University of São Paulo animal facility of the Department of Immunology - ICB. 8-week-old non-pregnant or 11-week-old pregnant female mice were used. All animals were maintained in accordance with institutional guidelines for animal welfare after approval by the Institutional Animal Care and Use Committee at University of São Paulo, as described above.

### ZIKV real-time quantitative RT-PCR

Viral RA from blood, serum or urine samples was extracted using the easyMAG mated extractor (BioMerieux, Quebec, Canada) or QIAamp viral RNA mini kit (Qiagen, Hilden, USA), according to manufacturer’s instructions. Estimation of viral RNA copy number in patients’ samples was performed using real-time RT-PCR (TaqMan RNA to-Ct 1-Step Kit; Applied Biosystems) with primers and probes, as previously described [9, 65]. qRT-PCRs with cycle threshold (Ct) values higher than 40 cycles were considered negative. Quantitative assay was performed using a standard curve produced with serial 10- fold dilutions of ZIKV RNA expressed on a log10 scale as genome equivalents/sample.

### Viral RNA Sequencing and Assembly

To evaluate the quantity and quality of the RNA extracted, a Qubit® 2.0 Fluorometer (Invitrogen, Carlsbad, USA) and an Agilent 2100 Bioanalyzer (Agilent Technologies, Santa Clara, USA) were used, respectively. Synthesis of cDNA was performed using SuperScript II and random hexamer primers according to the manufacturer’s recommendations (Invitrogen, Carlsbad, USA). Nucleotide sequencing was performed using the TruSeq RNA sample preparation kit in an Illumina HiSeq 2500 instrument (Illumina, San Diego, USA) with a paired-end and 150-base-read protocol in *RAPID* module. The sequencing reads were assembled *de novo* using the metaViC pipeline (available on https://github.com/sejmodha/MetaViC), as previously described [69].

### Viral quantitation by focus forming unit assay and plaque forming unit assay

The viral titter of viral isolates was determined by both focus forming unit (FFU) and plaque forming unit (PFU) assays in Vero E6 cells. The quantitation of viral load during in vitro experiments were performed by FFU assay, while the viral load in different tissues of animals were determined by PFU assay. For FFU quantification, samples were clarified by centrifugation (2,000 x g at 4°C for 10 min) and diluted serially prior to infection. These dilutions were added to Vero E6 cells in 96-well plates for 2 h for viral adsorption and, after supernatants removal, cells were maintained with MEM + 1% CMC medium (final concentration) supplemented with 5% FBS and 1% penicillin/streptomycin for 48 hours at 37°C and 5% CO_2._ The overlay was removed, and cells were fixed overnight with 1% paraformaldehyde (PFA) in PBS. To detect infected cell foci, cells were permeabilizated with Triton buffer (PBS 0.15M, 0.1% BSA and 0.1% Triton X-100) and foci were detected after incubation with a mouse anti-ZIKV NS1 antibody in a volume of 50 μl for 2 h at room temperature. After three washes with 300 μl of permeabilization-wash buffer (P-W; PBS, 0.1% saponin, and 0.1% bovine serum albumin [BSA]), the samples were incubated with 50 μl of a 1:2,000 dilution of horseradish peroxidase (HRP)-conjugated goat anti-mouse IgG for 1 h at room temperature. Focci were stained by the addition of the TrueBlue detection reagent (KPL), and the blue spots were counted after three washes with distilled water.

For PFU quantification, Vero E6 cells (1 x 10^5^/well) distributed in 24-well plate were culture with clarified samples from infected mice. These samles were diluted (10^5^ – 10^1^) in RPMI-16-40 medium and used to infect VeroE6 cells for 1h at 37ᵒC. Further, supernatants were removed and DMEM medium (2% FBS + 1% penicillin/streptomycin) + 1.5% CMC (Carboxymethyl cellulose) were added in each well. Cells were cultured in 37ᵒC for 5 days and then stained with crystal violet for plaque counting. Viral titter of the stock sample was determined by: [average number of plaques/ (dilution factor of well)] x (volume of inoculum per plate).

### ZIKV infection and treatment

For *in vitro* infection, PBMCs, THP-1 monocytes and hBMECs were infected with ZIKV MOIs (multiplicity of infection) ranging from 0.1 to 10, depending on the experimental design. In some experiments, cells were submitted to starvation (serum free medium) from 4 to 18 hours prior to infection with the virus for 2 h at 37°C. Afterwards, cells were washed four times with PBS. Culture medium was added to each well and the cells were incubated at 37°C and 5% CO_2_ for the duration of the experiment. For Gas6 γ- carboxylation and Axl kinase blockade experiments, cells were incubated or not (mock) with 2μM warfarin or 2μM R428 (BerGenBio), respectively. Cells were harvested after 1-, 2-, 3-, 4-, 6-, 9-, 12-, 24-, 48- or 72-hours post infection depending on the experimental design. Medium was harvested at specified time points for determination of Gas6 production, ZIKV titters and replication either by PFU or flow cytometry.

### Mice infection

For *in vivo* experiments in mice, the BeH815744 virus was incubated or not with 1µg/mL of recombinant mouse Gas6 (rmGas6) in μl for 4 hours at 37ᵒC prior to mice inoculation [20]. 8-week-old SJL or C57BL/6 WT mice were inoculated with 10^2^ ZIKV pfu diluted in 100μL of PBS or pre-incubated with 1μg/mL of recombinant murine Gas6 (ZIKV^rmGas6^) by intravenous route (retroorbital sinus). Mice were euthanized at days 1, 3 or 5 post infection depending on the experimental design. To study congenital infection, 11-week-old C57BL/6 WT pregnant mice were infected with ZIKV 10^5^ pfu in 30μL of PBS pre-incubated or not with 1μg/mL rmGas6 at embryonic day E10 or E16 intravaginally or subcutaneously (footpad), respectively. Tissues were harvested at E18 or postnatally (P) at day 4.

### Relative quantitation by Real-time quantitative RT-PCR

For *in vitro* analysis, RNA was extracted using the miRVana miRNA Extraction kit (Ambion) according to the manufacturer’s instructions. RNA quantity was determined by NanoDrop spectrometric dosing. Total RNA samples (up to 1 μg) were reverse transcribed using the oligo(dT) primer from the High Capacity cDNA Reversion Transcription Kit (Thermo Fisher Scientific, Waltham, MA, USA). mRNA expression of the TAM receptors Tyro-3 (*Tyro-3*), Axl (*Axl*) and Mer (*Mer*), Suppressor of cytokine signalling-1/2 (*SOCS1*), Interferon-β (*IFN-β*), Interferon-induced protein with tetratricopeptide repeats-1 (*IFIT-1*), Gas6 (*Gas6*), Gamma-glutamyl carboxylase (*GGCX*), Vitamin K Epoxide Reductase Complex subunit 1 (*VKORC1*) and the housekeeping gene Glyceraldehyde 3-phosphate dehydrogenase (*GAPDH*) were determined by qRT-PCR, using iTaq Universal SYBR Green Supermix (Bio-Rad, Hercules, CA, USA). Cycling conditions were the following: 2 minutes at 95 °C and 35 cycles of 15 seconds at 95 °C and 60°C for 1 minute; 31 seconds at 65 °C and 60 cycles of 65 °C for 5 s (+0.5 °C/cycle; ramp 0.5 °C/s). The oligonucleotides used are described in Supplemental Table S1. Cycling conditions were the following: 95 °C for 5 min; 45 cycles of 95 °C for 15 s, and 60 °C for 1 min. The median cycle threshold (C_t_) value and 2^-⊗⊗Ct^ method were used for relative quantification analysis and all C_t_ values were normalized to GAPDH. Results were expressed as means and SEM of biologic replicates. All qRT-PCR assays were performed on the CFX96 Touch™ Real- Time PCR Detection System (BioRad, Hercules, USA).

For *in vivo* analysis, RNA extraction from mice organs was performed according to the protocol provided by manufacturer TRIzol™ Reagent (Cat.No. 15596026). From the obtained RNA, we performed qRT-PCR according to the protocol provided by the manufacturer (Applied Biosystems Cat. No 4368813). The PCR reactions were done using TaqMan® gene expression assay with the following cycles 95°C for 2 minutes and 45 cycles of 95°C for 15 seconds and 60°C for 1 minute. Amplification was normalized by beta actin expression. Gene expression was analyzed by 2^−ΔΔCt^ method. Results were expressed as means and SEM of biologic replicates.

### Flow Cytometry

Vero E6 cells were incubated for 4 days with 72h p.i. supernatants from THP-1 monocytes infected *in vitro* with ZIKV MOI 1, submitted or not to the different pharmacological treatments (mock, pre- or 2h p.i. treatment with 2uM warfarin). Cells were fixed/permeabilized with Cytofix/Cytoperm (BD Biosciences, Franklin Lakes, New Jersey, U.S.) according to manufacturer’s instructions and incubated with 1:100 mouse anti-ZIKV NS1 antibody at 4°C for 30 minutes. Further, cells were washed with Permwash (BD Biosciences) and incubated with 1:100 Alexa Fluor 488-conjugated goat anti-mouse IgG H&L, washed and subjected to flow cytometry using a FACS Calibur (BD Biosciences). Flow data was analyzed with FlowJo V10.

### ELISA for Gas6 quantification

To measure Growth Arrest-Specific Protein 6 (Gas6) in patients’ serum samples and supernatants from cells *in vitro*, ELISA was performed using Human Gas6 DuoSet ELISA kit according to the manufacturer’s instructions (R&D Systems, Minneapolis, Minnesota, USA). Briefly, 96-well microplate was coated with the goat anti-human Gas6 capture antibody diluted to the working concentration in PBS and incubated overnight at room temperature. The plate was washed and incubated with blocking buffer (reagent diluent) at room temperature for 1h After washing, samples were added (dilution 1:50 for serum samples and dilution 1:25 for cell supernatant samples) and incubated for 2h. Next, plates were washed and biotinylated goat anti-human Gas6 detection antibody, diluted in blocking buffer was incubated for 2h at room temperature, followed by streptavidin-HRP incubation for 20 minutes at room temperature. Plates were washed and incubated for 20 min with substrate solution. Optical density was determined at 450 nm followed by wavelength correction at 540 nm.

### Single-nucleotide polymorphism of Gas6 (SNP)

Total DNA from patients’ peripheral blood was extracted using QIAmp DNABlood Mini Kit (Qiagen, Germantown, Maryland, USA) following manufacture’s instruction. Extracted DNA was used to perform PCR and electrophoresis analysis of the GAS6 Intron 8 c.834+7G>A SNP, as previously described [21]. Reactions were performed using Taq DNA Polymerase Recombinant kit (Thermo Fisher Scientific, Waltham, Massachusetts, USA) following manufacture’s recommendations. Thermocycling was as follows: 5 minutes at 94°C, 35 cycles of 30 seconds at 94°C, 30 seconds at 55°C and 45 seconds at 72°C, then followed by 5 minutes at 72°C and 4°C until digestion. Afterwards, amplicons were digested with restriction enzyme HhaI (New England Biolabs, Ipswich, Massachusetts, USA) at 37°C for 4 hours and separated in 1.5% agarose gel with ethidium bromide at 80V for 90 minutes. UV documentation in Gel Doc XR+ System (BioRad, Hercules, Califórnia, USA).

### Correlation coefficients and network analysis

Pearson’s or Spearman’s Rank correlation coefficients were determined to assess the association between Gas6 and ZIKV-specific immune signatures determined in the serum samples from healthy donors and ZIKV-infected patients by Multiplex Microbead-Based Immunoassay, as previously described [9]. Correlation coefficients were also determined to assess the association between Gas6 levels in the serum and gene expression in the matched peripheral blood cells from ZIKV-infected adult patients. Correlations between transcripts were also quantified in the cells infected *in vitro* with ZIKV. For the network analysis, the systemic level of each biomarker was an input in the R software (v. 3.4.3). Along with the rank-order correlation coefficient, the p-value to test for non- correlation was evaluated using p < 0.05 as a cutoff. Next, correlation networks were generated by the analysis of relationships among each biomarker in the serum samples. Based on the correlation coefficients, the same software was applied to identify links (edges) of interactions between the biomarkers (nodes). The correlation strength is represented by edge transparency and width; positive correlations are represented by red edges, and negatives correlations are represented by blue edges. Following this approach, each biomarker was selected as a target, and the R software was used to perform a search within the other biomarkers for those that were associated with the target, in terms of correlation strength. As a result, the features related to the selected target were linked. This process was repeated for each biomarker, and the result was the inferred network among the input values. To analyze the structure of the networks, the graphs for the network analysis were customized in the Cytoscape software (v 3.5.1) using the pre-fuse force-directed layout. This layout follows an algorithm that in the equilibrium state for the system of forces, determined by the correlation strength, the edges tend to have uniform length, and nodes that are not connected by an edge tend to be drawn further apart

### Statistical Analysis

Data normality was checked by the Shapiro-Wilk test. Two-tailed Student’s t-test or One-way analyses of variance statistical test with Bonferroni-corrected multiple comparisons test were used to compare means between groups with normally distributed data. Data sets with non-parametric distributions were compared using the Mann– Whitney test or Kruskal-Wallis test with post hoc correction for multiple testing using the original FDR method of Benjamini and Hochberg, with p < 0.05 considered significant. Data are presented as Tukey box plots or means and SEM, unless otherwise stated, of 2–3 representative and independent experiments with similar results. Analysis were performed and the graphs generated in GraphPad Prism 8 and R software. p<0,05 were considered relevant.

## NOTES

### Author contributions

Conceptualization, J.L.S.F, L.G.O, F.T.M.C., J.P.S.P. and J.L.P.-M.; Methodology, J.L.S.F., L.G.O, L.M., P.L.P., N.G.Z., C.M.P., C.L.F., D.A.T.-T., W.M.S., M.R.A., J.F., S.P.M., G.F.S., C.C.J.; Investigation, J.L.S.F., L.G.O, L.M., P.L.P., N.G.Z., C.M.P., D.A.T.-T., W.M.S., M.R.A., J.F., S.P.M., G.F.S., C.C.J., L.F.P.N,; Clinical Data Acquisition, M.C.M., K.B.S., R.N.A., A.R.R.F.; Patient enrolling, M.L.C., R.N.A., A.R.R.F., M.R.R., M.T.G., M.L.M.; Original Draft, J.L.S.F. and L.G.O; Review & Editing, W.M.S., L.F.P.N., C.V.R, F.T.M.C., J.P.S.O., J.L.P.-M.; Visualization, L.M., P.L.P., N.G.Z., C.M.P., D.A.T.-T., W.M.S., M.R.A., J.F., S.P.M., G.F.S., C.C.J., M.C.M., K.B.S., M.L.C., R.N.A., A.R.R.F., M.R.R., M.T.G., M.L.M; Funding Acquisition, F.T.M.C, J.P.S.P. and J.L.P.-M.; Resources, L.F.P.N., C.V.R., F.T.M.C, J.P.S.P., J.L.P.-M.; Supervision, F.T.M.C, J.P.S.P. and J.L.P.-M.

### Acknowledgments

We thank the study participants and healthy volunteers for their participation and the clinical staffs from University of Campinas Hospitals and other hospitals of this city for assistance in patient enrolment and healthcare, blood sample preparation, study coordination, and data entry. We thank the staff of the Life Sciences Core Facility (LaCTAD) from Unicamp for the High-throughput sequencing.

### Disclaimer

The funders had no role in study design, data collection and analysis, decision to publish, or preparation of the manuscript.

## SUPPLEMENTAL FIGURES

**Supplementary figure 1:**
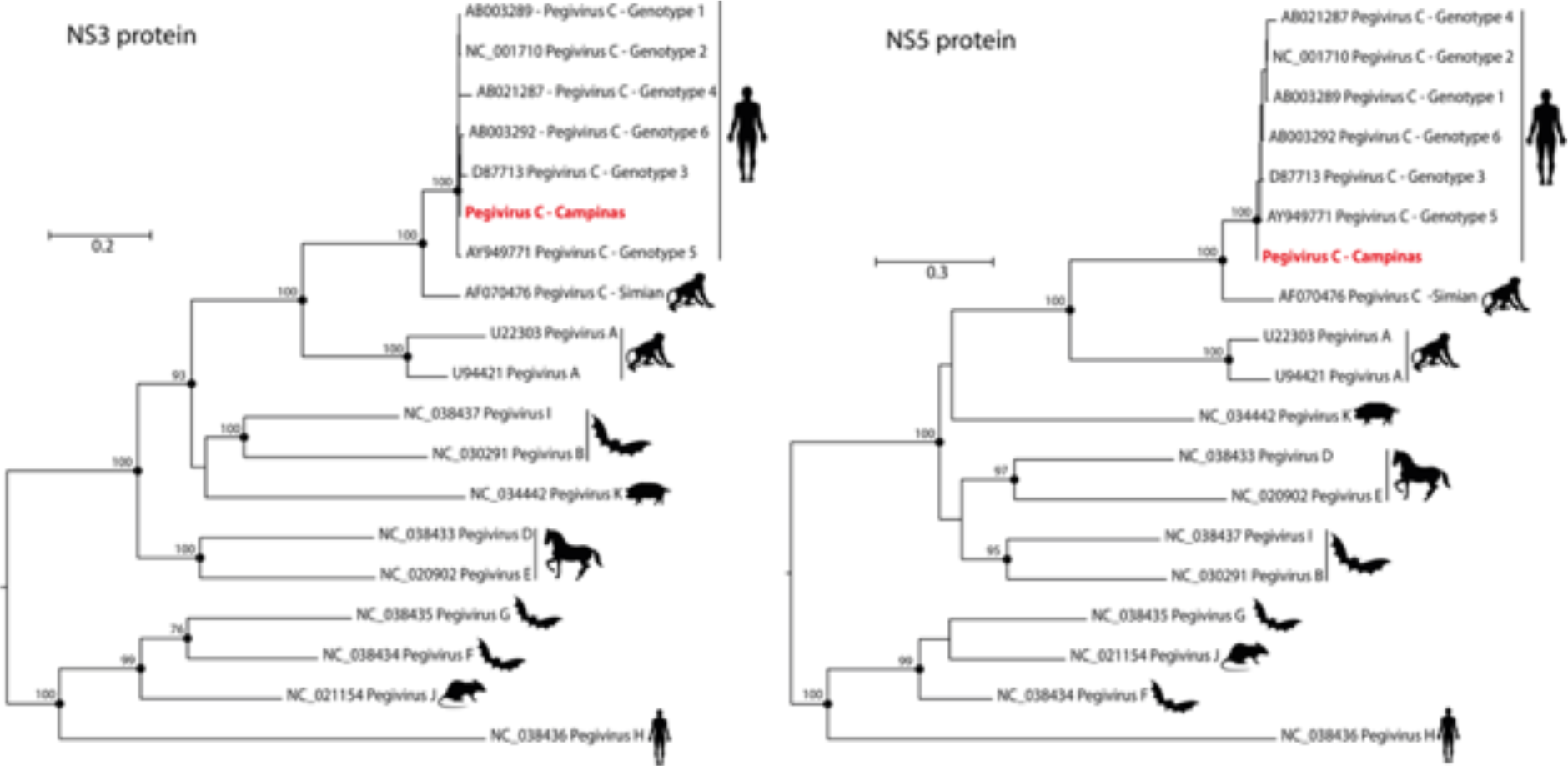
The virus detected by high-throughput sequencing in the Neuro^ZIKV^ patient is a human pegivirus. Phylogenetic trees of maximum likelihood showing the evolutionary relationships of the detected human pegivirus (named of human pegivirus strain Campinas, highlighted in red) with representative members of the *Pegivirus* genus according to the alignment of the NS3 and NS5 genes. Bootstrap values are indicated inside black circles, and hosts are indicated by figures.

**Supplementary figure 2:**
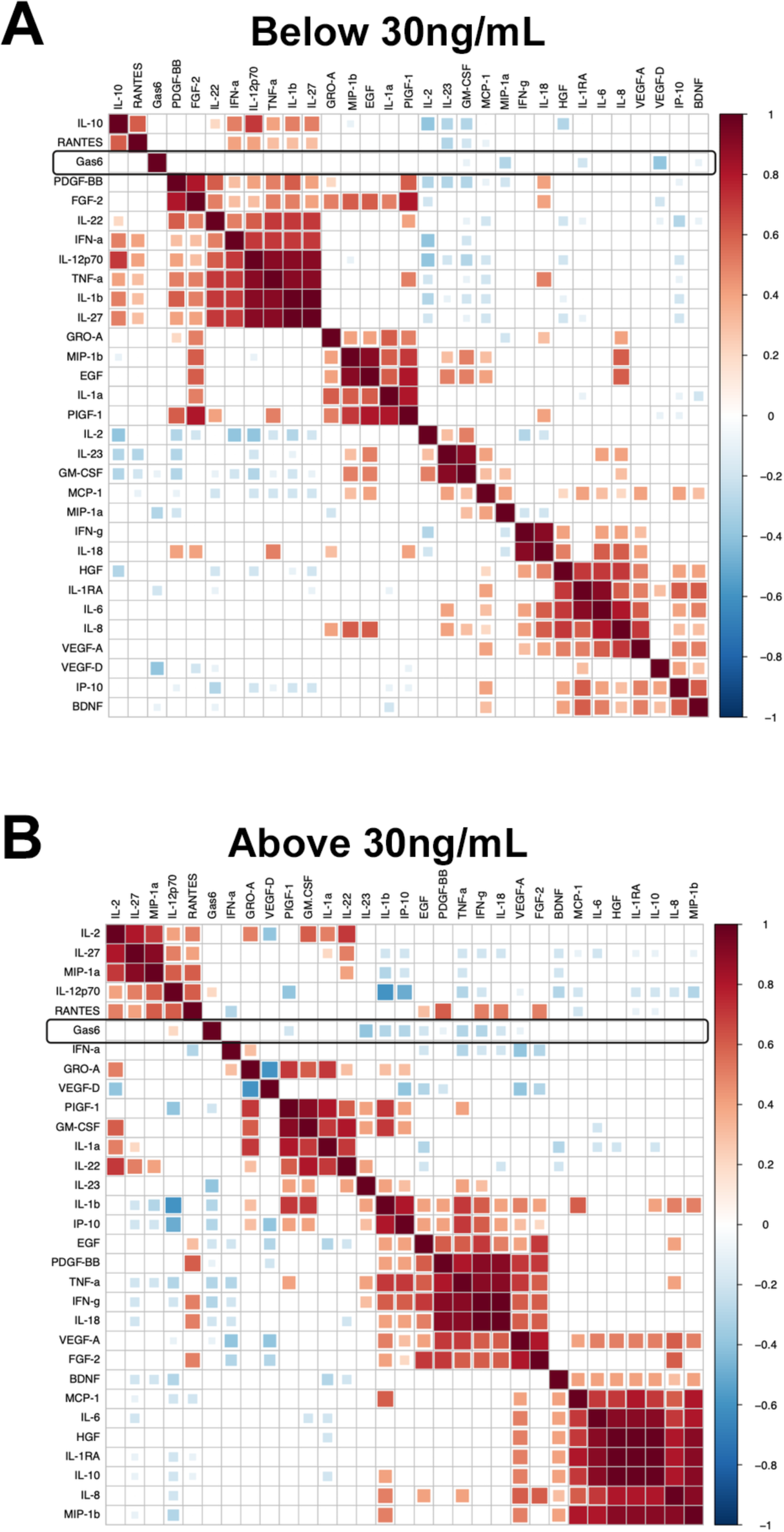
Estimation of Gas6 threshold changing correlations with ZIKV-specific immune signatures. A reduced complexity model was established by focusing on informative interactions between ZIKV- specific immune signatures and Gas6 determined by Pearson’s correlation coefficients based on Gas6 levels. (A) Below 30ng/mL; (B) Above 30ng/mL Only correlations with associated *p*-value <0.05 are shown.

**Supplementary figure 3:**
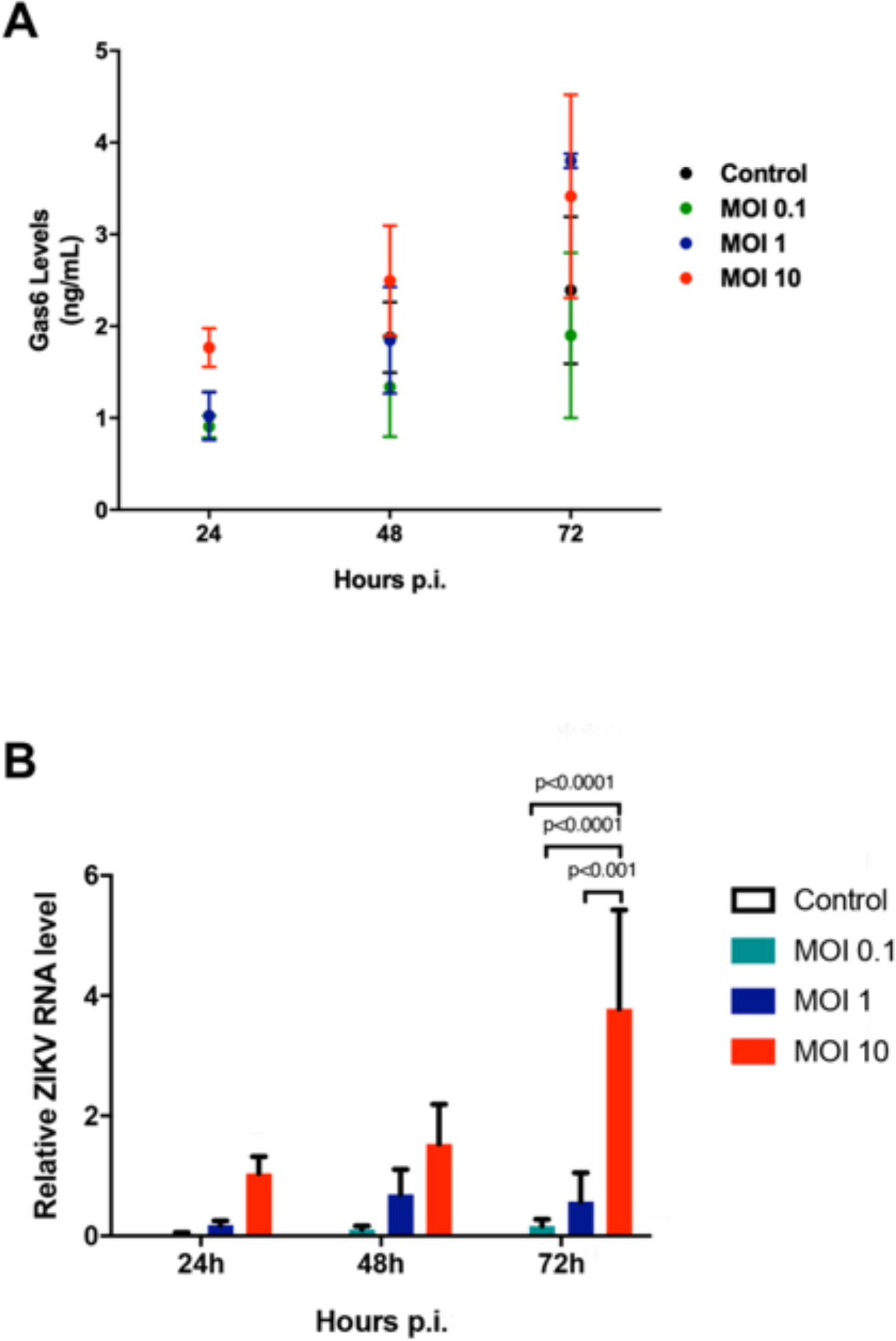
Gas6 production by human brain microvascular endothelial cells (hBMECs) *in vitro* is not affected by ZIKV infection. (A) Gas6 levels in the supernatant of hBMEC at different time points (24, 48 and 72h) after *in vitro* infection with ZIKV at different multiplicities of infections (MOI 0.1, 1 and 10). (B) Total cellular RNA was extracted at different time points after infection as indicated in the graphs, and relative viral RNA levels were determined by real-time quantitative PCR. The data shown are representative images of 3 independent experiments. P values were determined using one-way ANOVA statistical test with for each time point with Bonferroni-corrected multiple comparisons test.

**Supplementary figure 4:**
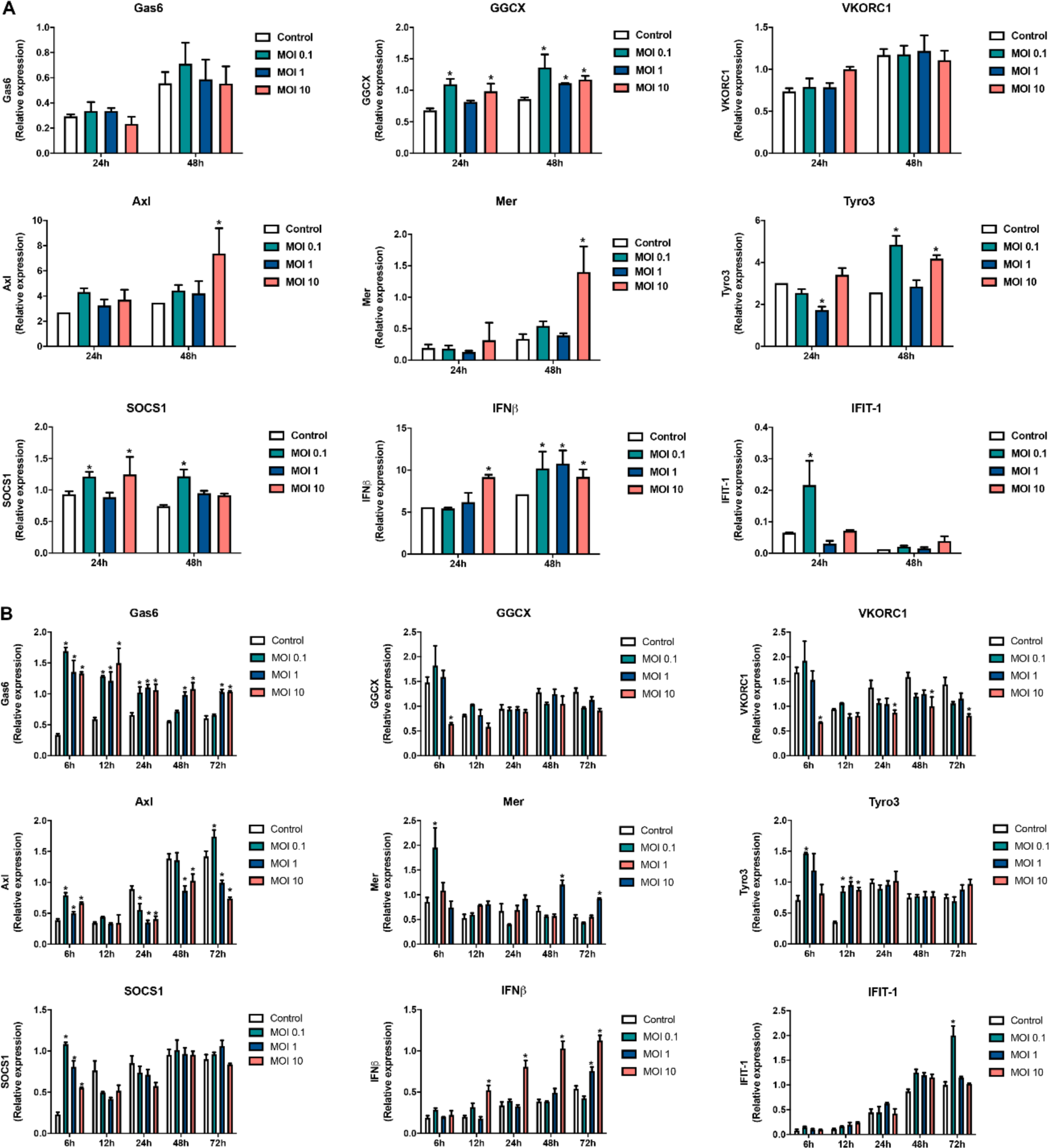
ZIKV Infection stimulates Gas6 upregulation and downregulates type I IFN response in peripheral blood mononuclear cells (PBMCs) and monocytes. (A) PBMCs and (B) THP-1 monocytes were challenged with ZIKV (MOI 0.1, 1 and 10), respectively. Total cellular RNA was extracted at different time points after infection as indicated in the graphs, and relative viral RNA levels, *GAS6*, *GGCX*, *VKORC1*, *AXL*, *TYRO3*, *MER*, *SOCS1*, *IFNB* AND *IFIT1* mRNA levels were determined by real-time quantitative PCR. (A) The data shown are mean ± SEM representative of three independent experiments using cells from different donors. P values were determined for comparisons between conditions at the corresponding time point using Two-way analyses of variance statistical test with Tukey-corrected multiple comparisons test. \**p* <0 .05 vs control (uninfected cels).

**Supplementary figure 5:**
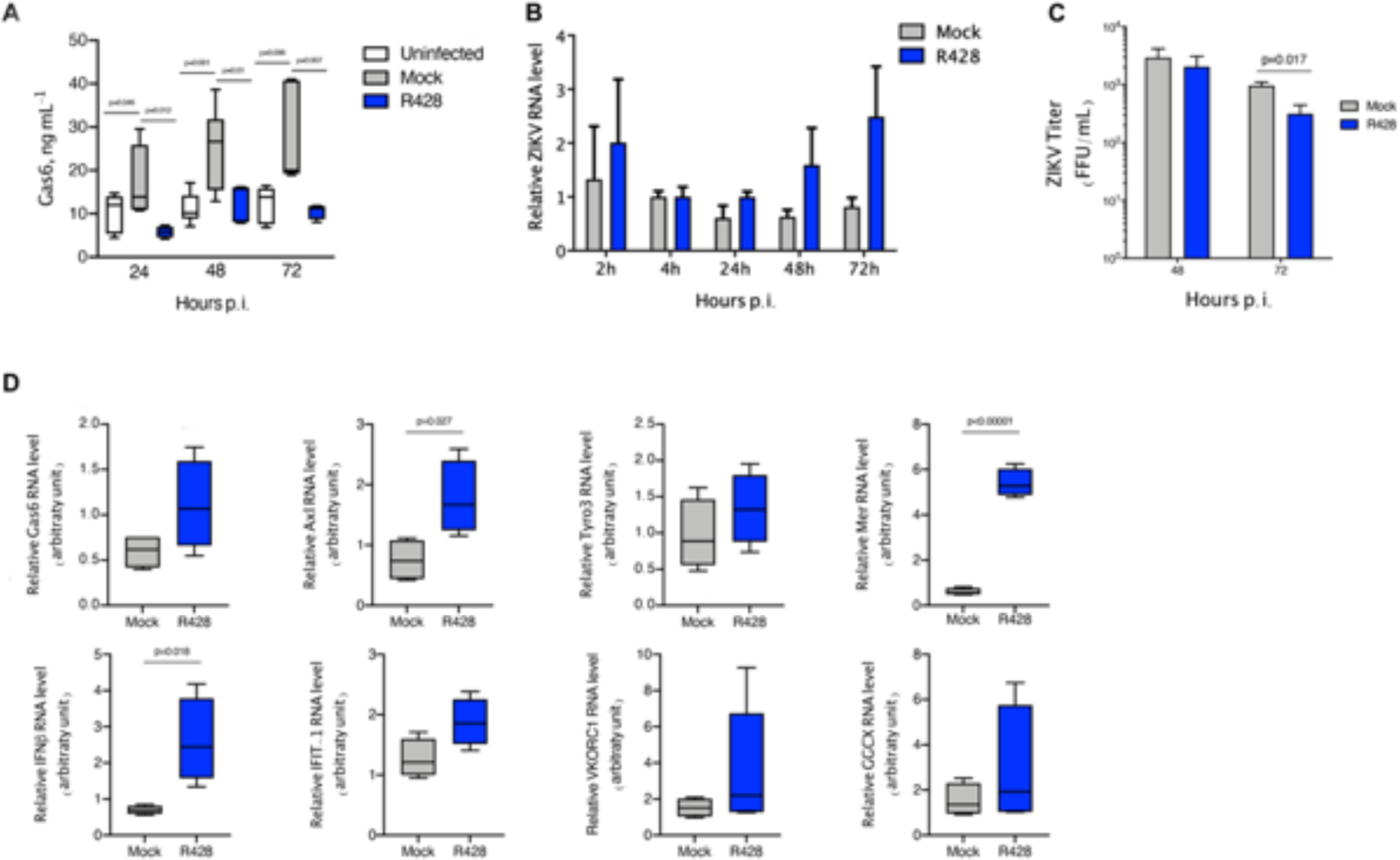
R428 treatment, inhibitor of Axl tyrosine kinase activity, restores antiviral response. (A) Gas6 levels were determined by ELISA in the supernatant of THP-1 monocytes collected at 24h, 48h and 72h after *in vitro* infection with ZIKV (MOI 1). Infected cells were treated or not (mock) with 2µM R428 throughout the course of infection. Gas6 concentration is depicted as Tukey box plots. The data shown are representative images of three independent experiments. P values were calculated for comparisons between conditions at the corresponding time point using One-way analyses of variance statistical test with Bonferroni-corrected multiple comparisons test. (B, D) THP-1 monocytes were challenged *in vitro* with ZIKV (MOI 1) and treated or not (mock) with 2µM R428 throughout the course of infection. Total cellular RNA was extracted at different time points after infection. Relative viral RNA levels, *GAS6*, *GGCX*, *VKORC1*, *AXL*, *TYRO3*, *MER*, *SOCS1*, *IFNB* AND *IFIT1* mRNA levels were determined by real-time quantitative PCR. The data shown are representative images of three independent experiments. (B) The data shown are mean ± SEM representative of three independent experiments. (D) Graphs depict relative RNA expression as Tukey box plots after results were normalized to GAPDH housekeeping gene expression. P values were calculated using student’s t-test to compare groups with normally distributed data or Mann-Whitney test to compare groups with non-normal distributions. (C, D) ZIKV titer (FFU/mL) was determined by plaque forming assay after incubation of ZIKV-permissive Vero E6 cell line with the 48h or 72h supernatant from monocytes after infection *in vitro* with ZIKV (MOI 1), treated or not (mock) with 2µM R428 at the moment of infection (pre-treatment). The data shown are representative images of three independent experiments. Mann-Whitney test was used to compare conditions at the corresponding time point.

**Supplementary figure 6:**
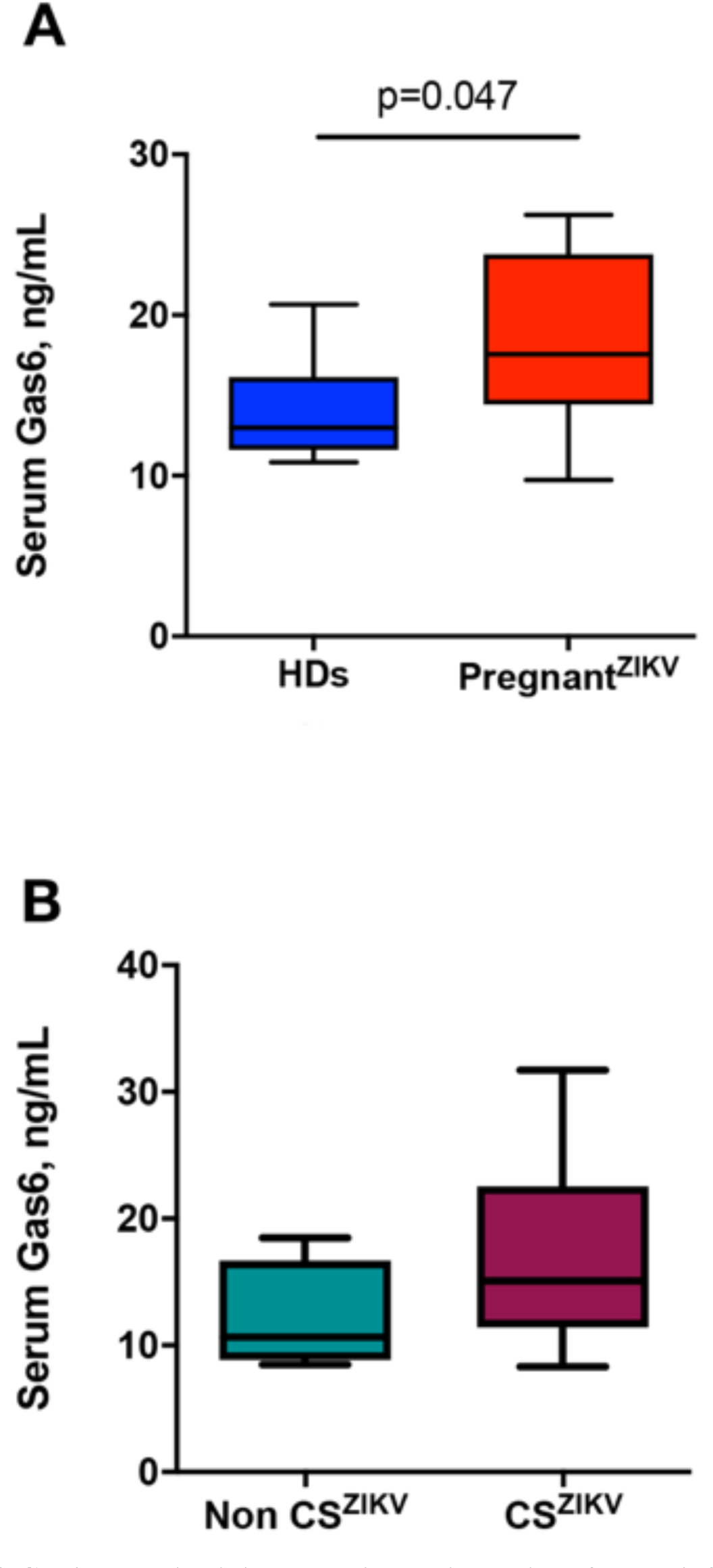
Gas6 expression is increased in the circulation of ZIKV-infected pregnant women. (A) Levels of Gas6 in acute-phase serum samples of 8 Zika virus (ZIKV)- infected pregnant women and 6 infants born to women with Zika virus (ZIKV) infection, were determined by ELISA. (B) Two infants were born with CNS abnormalities associated to ZIKV congenital syndrome (CS^ZIKV^) and 4 without congenital syndrome (Non CS^ZIKV^). Gas6 concentration is depicted as Tukey box plots. P values were determined using student’s t-test.

**Supplementary figure 7:**
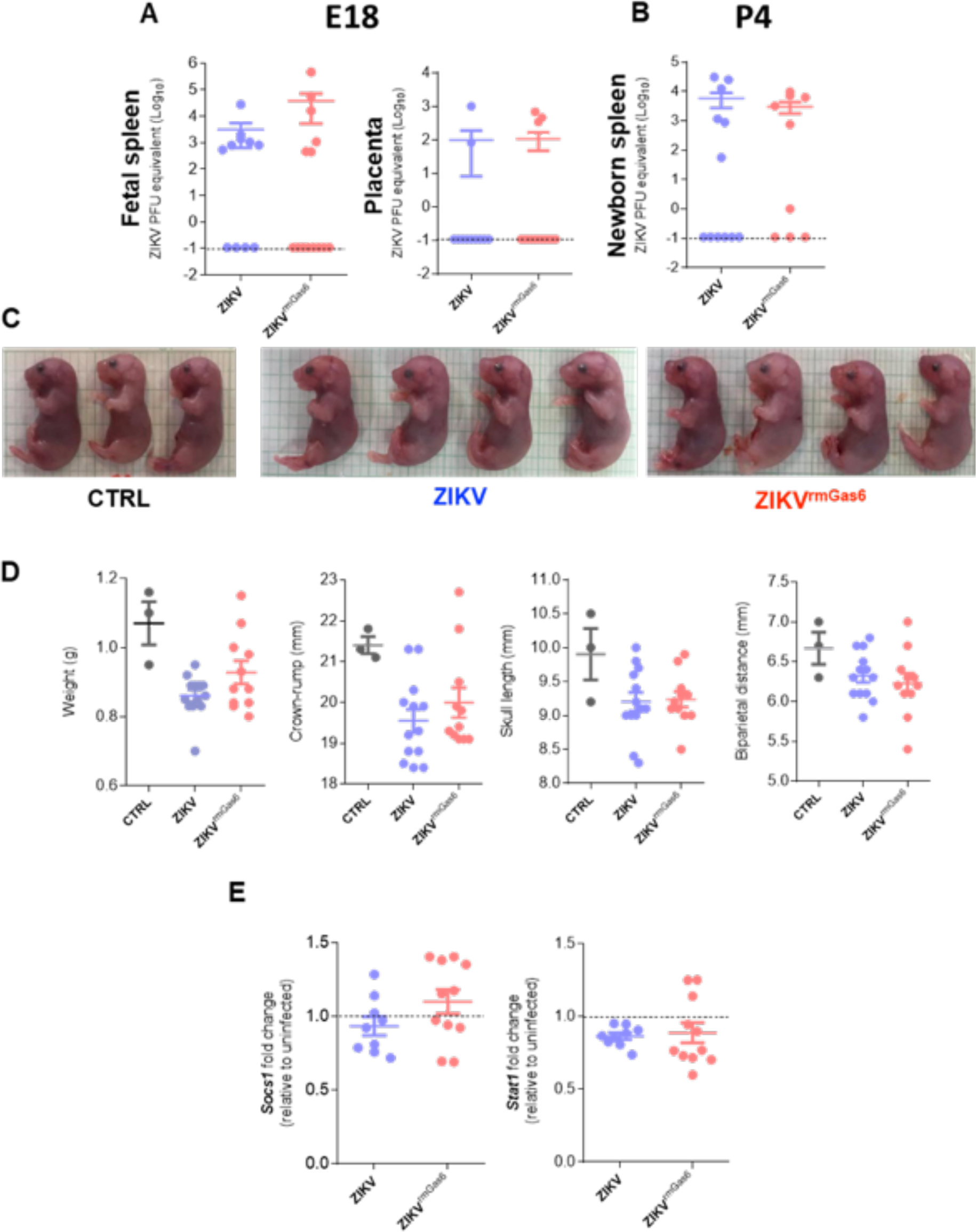
Dimension analysis and ZIKV viral load in C57BL/6 offspring. Pregnant mice were infected subcutaneously with pure ZIKV (10^5^) or ZIKV previously incubated with rmGas6 (1ug/mL)) (ZIKV^rmGas6^) on E (embryonic day) 16. The organs were harvested on E18 or postpartum day (P) 4. The viral load was analyzed by qPCR. Viral load on E18 (A) and P4 (B). Pregnant mice were also infected intravaginally on E10 with pure ZIKV (10^5^) or ZIKV previously incubated with rmGas6 (1ug/mL). Tissues were harvested at E18. Foetus pictures (C). Dimension analyses (D). Placenta gene expression (E). Fold change was calculated between uninfected and infected groups. Graph bars are shown as mean ± SEM and are representative of two independent experiments. Numbers of experimental groups: (A) Control n=6 ZIKV n=11; ZIKV^rmGas6^ n=14. (B) Control n=6 ZIKV n=12; ZIKV^rmGas6^ n=9. (C and D) Control n=3 ZIKV n=13; ZIKV^rmGas6^ n=11. (E) Control n=3; ZIKV n=9; ZIKV^rmGas6^ n=11.

